# E-cigarettes compromise the gut barrier and trigger gut inflammation

**DOI:** 10.1101/2020.07.29.227348

**Authors:** Aditi Sharma, Jasper Lee, Ayden G. Fonseca, Alex Moshensky, Taha Kothari, Ibrahim M. Sayed, Stella-Rita Ibeawuchi, Rama F. Pranadinata, Jason Ear, Debashis Sahoo, Laura E. Crotty-Alexander, Pradipta Ghosh, Soumita Das

**Affiliations:** Department of Pathology, University of California, San Diego, CA, 92093; Department of Cellular and Molecular Medicine, University of California, San Diego, CA, 92093; Department of Medicine, University of California, San Diego, CA, 92093; Veterans Affairs Medical Center, La Jolla, San Diego, CA, 92093; Rebecca and John Moore Comprehensive Cancer Center, University of California, San Diego, CA, 92093

## Abstract

E-cigarette and vaping device use continue to rise, particularly in adolescents and young adults, but the safety of inhaling the multitude of chemicals within e-cigarette aerosols has been questioned. While several studies have evaluated vaping effects on the lungs and heart; effects on the gastrointestinal tract remain unknown. Using established murine models of acute (1 week) and chronic (3 month) daily e-cigarette aerosol inhalation, both with nicotine-containing and vehicle control e-liquids, murine colon transcriptomics and organoid co-culture models, we assessed the effects of e-cigarette use on the gut barrier and mucosal health. Histologic analyses revealed that chronic exposure to nicotine-free e-cigarette aerosols induced mucosal inflammation. Transcriptome analyses revealed that chronic, but not acute, nicotine-free e-cigarette use significantly reduced expression of tight junction markers, including occluding, and drove expression of pro-inflammatory cytokines. Exposure of murine and human enteroid-derived monolayers (EDMs) to nicotine-free e-cigarette aerosols alone, or in co-culture with invasive *E. coli,* confirmed that repetitive exposure was sufficient to recapitulate the key findings observed *in vivo*, i.e., barrier-disruption, downregulation of occludin, inflammation, and an accentuated risk of and response to bacterial infection. These data highlight an unexpected harmful effect of e-cigarette use on the gut barrier and pinpoint non-nicotine chemical components common across >90% of e-cigarette e-liquids as the source of harm. Given the ever-expanding importance of the integrity of the gut barrier for host fitness, and impact of gut mucosal inflammation on a multitude of chronic diseases, these findings are broadly relevant to medicine and public health.

**SIGNIFICANCE:** The safety of electronic cigarettes has been questioned amidst emerging evidence that they may derail our immune system and increase our susceptibility to infections. Despite these insights, their impact on the most critical entity that separates trillions of microbes from the largest immune system in our body, i.e., the gut barrier, remains unexplored. Using a combination of mouse models, gut transcriptomics, and murine and human gut-derived organoids, here we show that chronic exposure to aerosols of electronic-cigarette disrupts the gut barrier, increases its susceptibility to bacterial infections and triggers inflammation. Given the importance of the gut barrier in the maintenance of immune homeostasis, these findings provide valuable insights into the potential long-term harmful effects of electronic cigarettes on health.

## INTRODUCTION

Electronic nicotine delivery systems (ENDS), commonly referred to as e-cigarettes and vaping devices, were introduced to the international market in 2007 (1). Since then, e-cigarettes have become widely popular in the United States (2), primarily among the nation’s youth. Rapid growth in consumption is fueled in large part due to successful social media-based marketing and availability of appealing flavors and sleek devices. Although a large focus of research and regulation has been on the nicotine delivery function of these devices, because of the high addictiveness and known adverse health effects of nicotine, the majority of e-liquids contain propylene glycol (PG), glycerol (Gly), flavorants and contaminants, all of which may cause their own adverse effects on health (3–8). Regardless of chemicals contained, the popularity of e-cigarettes is driven, in part, by the notion that e-cigarettes represent a risk-free alternative to combustible cigarettes, and hence, few regulations exist to control the quality and composition of the ingredients used in e-cigarettes (7, 9).

This concept of risk-free use has recently been challenged (10). For instance, morbidity and mortality among teenagers and young adults due to the e-cigarette or vaping product use-associated lung injury (EVALI) epidemic (11) created a public health crisis in 2019-2020. This disease was found to be primarily associated with a single (non-nicotine) chemical included in vaping liquids: vitamin E acetate (12). Prior to the identification of EVALI, *in vitro* and *ex vivo* research studies had showed that e-liquids induce inflammatory responses and alter innate immune defenses in myeloid and primary airway epithelial cells (13–15). *In vivo* studies in which C57BL/6 mice were exposed to e-cigarette aerosols for 2 weeks found impairment of both pulmonary bacterial and viral clearance (16, 17), which raises alarm for increased susceptibility to influenza and coronavirus infections in particular. Evidence of pulmonary and systemic inflammation has also been found in the plasma (18) and bronchoalveolar lavage (19) samples from human e-cigarette-users, with elevated biomarkers of inflammation, e.g., (interleukin (IL)-1β, IL-6, IL-8, IL-13 and interferon (IFN)-γ. Furthermore, mechanisms that may connect the use of e-cigarettes and an increased risk for cancer development, as well as their stimulatory effect on cancer progression, have been proposed (20) and e-cigarettes have been shown to induce DNA damage [independent of nicotine(8)] while reducing repair pathways (21). Tang et al. determined that chronic e-cigarette use for one year led to increased development of lung cancer in mice (22). Recently, a NIH-funded panel reviewed the current evidence and came to the consensus that all chemical components of e-cigarette and vaping aerosols have the potential to cause distinct health effects (23), both similar to and disparate from those of nicotine or conventional tobacco smoke, both in the heart and lungs and throughout the body (23).

Here we set out to assess the impact of e-cigarette aerosol inhalation (with or without nicotine) on the gastrointestinal (GI) tract. The gut is resident to diverse microbiota (symbionts and pathobionts) and its handling of and response to the same is known to regulate several chronic diseases such as inflammatory bowel diseases (IBD), obesity, cardiovascular diseases, cancers and rheumatoid arthritis (24). Beyond the fact that e-cigarette use significantly modulates the oral microbiome by increasing the abundance of oral pathobionts (25), but is not known to significantly impact the gut microbiome (26), nothing is known as to how e-cigarette use may influence the gut barrier. Using a combination of mouse models, transcriptomics, and murine and human gut-derived organoids as *ex vivo* near-physiologic model systems, here we expose the hitherto unknown effects of e-cigarettes on the gastrointestinal tract and provide insights into the potential long-term effects of e-cigarettes on health.

## RESULTS

### Daily e-cigarette aerosol inhalation drives inflammation in the colon and reduces expression of genes related to barrier function

E-cigarettes function by heating liquid solutions and then pulling them through an atomizer to create an aerosol that is inhaled. To establish the *in vivo* effect of inhalation of these aerosols on the colon, the distal colon was harvested from mice exposed daily to e-cigarette aerosols at two time-points: 1 week (resembling acute exposure) and 3 months (resembling chronic exposure) (**Fig 1A**). Because the most common chemicals in e-cigarette aerosols are nicotine and humectants (propylene glycol and glycerol), we utilized nicotine-free and nicotine-containing (6 mg/ml) e-liquids with a 70:30 ratio of propylene glycol and glycerol within a Kanger Subtank attached to a box Mod e-device, and used exposure chambers with room air for controls (**Fig 1A**). H&E staining of distal colons from mice acutely exposed to nicotine-free (vehicle only) e-cigarette aerosols (e-cig) demonstrated small, infrequent patches of leukocyte infiltration in the submucosal layers (**Fig 1B**, *left; asterisk*). Acute exposure to nicotine containing aerosols (e-cig + nicotine) was associated with infrequent patches of epithelial erosions. By contrast, chronic e-cig exposure led to large submucosal inflammatory infiltrates within the colon (**Fig 1B**, *right*). No inflammatory infiltrates were seen in air controls, while smaller and infrequent infiltrates were present in colons of mice exposed to e-cig + nicotine.

**Figure 1:**
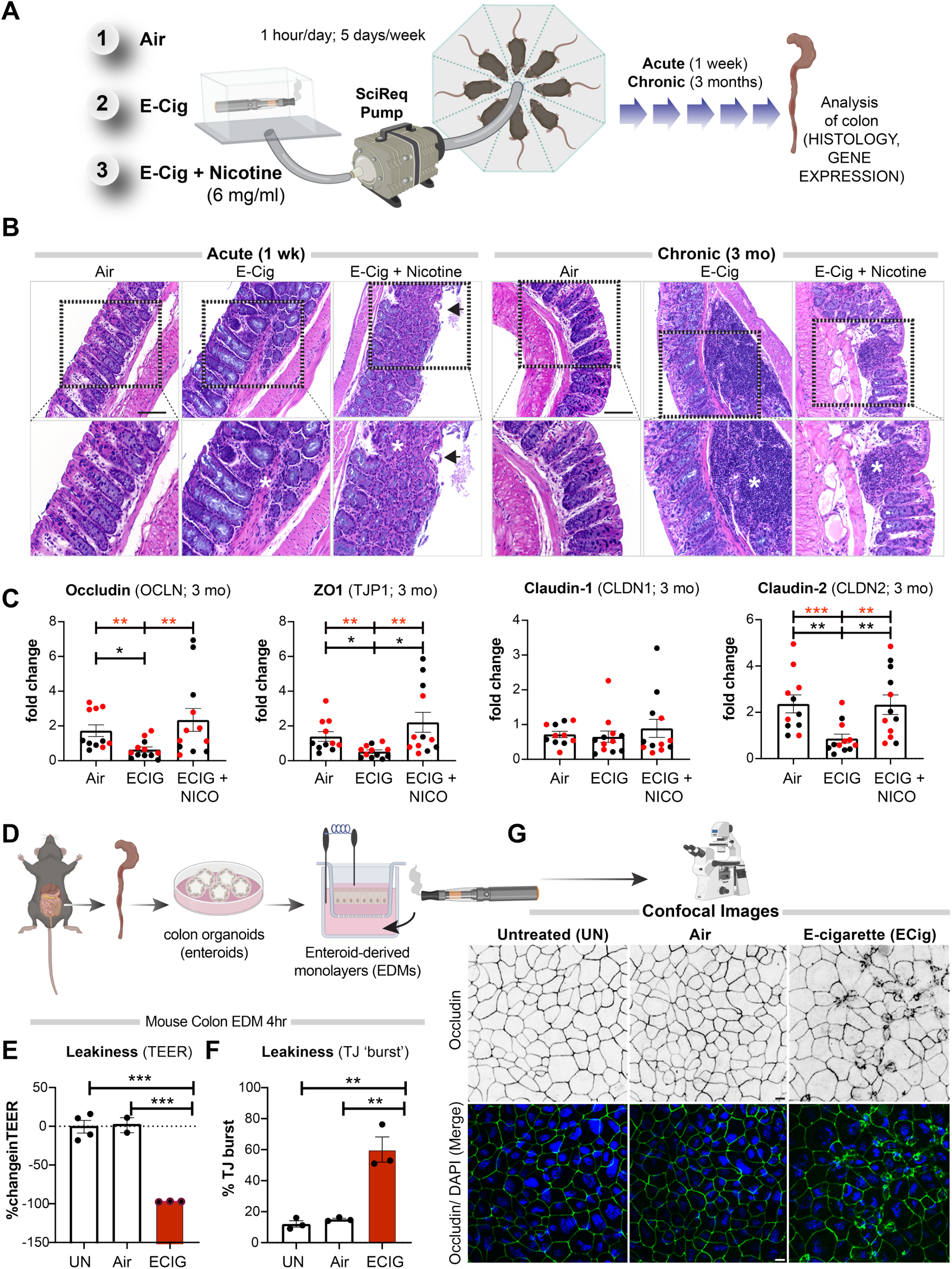
Electronic cigarettes trigger inflammation in the murine colon and disrupt the integrity of the murine gut barrier. **A**. Schematic displays the key aspects of the murine model of vaping used in this study. Mice were exposed either to air (negative control), or nicotine-free e-cig (e-cig alone; MOD brand) or nicotine-containing e-cig (e-cig + 6 mg/ml nicotine) for 1 wk or 3 mo. **B**. Hematoxylin-eosin staining of distal colons after 1 wk (left) or 3 mo of exposure to e-cig. Asterisks = inflammatory infiltrates; arrowheads = epithelial erosions. **C**. Bar graphs display the relative levels of expression of genes in the colon (black dots, male mice; red dots, female mice) that encode proteins that regulate epithelial tight junctions. Data is displayed as mean ± SEM. Statistical significance was estimated using either one-way ANOVA with Tukey’s test (black) or Mann-Whitney’s test (red); *p◻<◻0.05, **p <◻0.01 and ***p <◻0.001. **D.** Schematic displays the key aspects of *ex vivo* disease modeling to interrogate the impact of vaping on the murine colonic epithelial barrier. **E.** Bar graphs display the percent change in TEER. Data is displayed as mean ± SEM (n = 3-5 independent experiments). **F-G.** EDMs were treated as indicated, fixed and stained for occludin (green) and DAPI (blue, nuclei) and analyzed by confocal microscopy. Bar graphs in **F** display the % increase in the tight junction (TJ) ‘bursts’ (indicative of disrupted TJs). Data is displayed as mean ± SEM (n = 3-5 fields/condition). Statistical significance was estimated using one-way ANOVA with Tukey’s test; **p <◻0.01. Confocal microscopic images in **G** are representative of EDMs, either untreated (UN) or after 4 h of treatment with air or e-cig-infused media. Scale bar = 10 μm..

Markers of gut epithelial tight junctions, e.g., occludin (OCLN), zonula occludens (ZO)-1 (TJP1) and Claudin-2 (CLDN2) had significantly reduced gene expression in mice chronically exposed to nicotine-free aerosols (e-cig) compared to air controls (**Fig 1C**). The fact that e-cig exposure affects the levels of Claudin-2, a major regulator of TJ-specific obliteration of the intercellular space (27), but not its counterpart Claudin-1 (**Fig 1C**), which is specialized for TJ integrity in the skin epidermis indicates that the effects of e-cig on TJs may be gut specific. No reduction was observed in any of the barrier function genes in mice exposed to nicotine-containing e-cigarettes. No significant differences were observed in levels of pro-inflammatory cytokines MCP1 or IL-8 in the chronically exposed mice (**Fig S1**). Finally, no statistically significant differences were observed in the transcript levels of TJ markers and pro-inflammatory cytokines in acute exposures between any conditions (**Fig S1**).

These findings indicate that chronic, but not acute exposure to aerosols of nicotine-free e-cigarettes is sufficient to trigger inflammation in the gut and that such inflammation is associated with reduced expression of markers of epithelial TJs. Findings also suggest that concomitant exposure to nicotine may alleviate both phenotypes.

### E-cigarette use disrupts the integrity of the gut barrier

The integrity of the gut barrier is maintained by a complex cross-talk between the gut epithelial barrier and immune cells in the lamina propria. To determine if the observed decrease in colon TJ markers in mice exposed to e-cigs is a direct consequence of aerosols on the gut epithelial barrier, we used an *ex vivo* near-physiologic model system called the “gut-in-a-dish” (**Fig 1D**). In this model, crypt-derived stem cells isolated from mouse colon (see the Materials and Methods section) were used to generate organoids and later differentiated into polarized enteriod-derived monolayers (EDMs). These EDMs have been validated as model systems that closely resemble the physiologic gut lining in which all cell types (enterocytes, goblet, paneth enteroendocrine, and tuft cells) are proportionately represented (28–31). We exposed the basolateral surfaces of the murine colonic EDMs to e-cigarette aerosol-infused media (or air-infused media; as controls) and analyzed the integrity of the gut barrier using two readouts (**Fig 1D**): (i) paracellular permeability, as reflected by low trans-epithelial electrical resistance (TEER) and (ii) molecular characterization of epithelial TJs by looking at the localization of occludin; this integral membrane protein allows us to not just visualize but also quantify the degree of TJ disruption. Exposure to nicotine-free e-cigarette aerosol media caused a significant drop in TEER (~% change value of −97.3±0.3%) compared to untreated (UN; −0.5±8.1%) and air-treated (1.4±9.7%) controls (**Fig 1E**). Findings indicate a significant increase in paracellular permeability upon exposure to e-cig when compared to untreated (p = 0.0002) and air-treated (p = 0.0004) controls (**Fig 1E**). In e-cig exposed EDMs, confocal microscopy showed significantly increased “burst” tricellular TJs [these are specialized regions of the TJ where three or more cells come in contact (32), are also the regions where TJ-disruption can be visualized/assessed first (33)] by ~60.1±8.1% when compared to untreated (p = 0.0010) and air-treated (p = 0.0015) controls (**Fig 1F-G**). These findings show that chemicals contained within the most basic e-cigarette aerosols disrupt the epithelial barrier. Because these effects were seen in *in vitro* assays on isolated gut epithelial monolayers in the absence of immune cells, findings also suggest a direct cause and effect relationship between e-cigarette aerosol components and epithelial barrier disruption. Because the chemicals used to make the e-liquids and e-cig aerosols used in these studies (propylene glycol and glycerol) are found in >99% of all e-cigarettes, these data broadly apply to e-cigarettes and vaping devices.

### Chronic exposure to inhaled nicotine-free e-cigarette aerosols induces stress-responses in the colon

To determine the global impact of e-cigarettes on the gut, we next carried out RNA sequencing on the distal colons. While acute exposure to nicotine-free e-cigarettes did not significantly change gene expression in the colon (**Fig S2A**), chronic exposure was associated with significant changes (**Fig 2A**). A differential expression analysis showed that chronic exposure to nicotine-free e-cigarettes was associated with a significant upregulation of 120 genes and downregulation of 75 genes (with a 30% false discovery rate, FDR) (**Fig 2B; Table S1**). Barring a handful of genes (arrowheads, **Fig 2B, S2B**) these differences were virtually reversed when mice were exposed to nicotine-containing e-cigarettes (**Fig 2B**). TJ markers occludin and ZO1 were downregulated by nicotine-free, but not nicotine-containing e-cigarettes (**Fig 2C-D**). Multiple pro-inflammatory cytokines were either elevated significantly (MCP1, IL-8, TNFα) or showed an increasing trend but did not reach significance (Cxcl2) (**Fig 2E-H**). These RNA Seq findings are in agreement with our prior observations by histology (**Fig 1B**) and the targeted analyses of TJ markers by qPCR (**Fig 1C**), in that, colons of mice exposed to nicotine-free, but not nicotine-containing e-cig have impaired tight junction markers and are inflamed.

**Figure 2:**
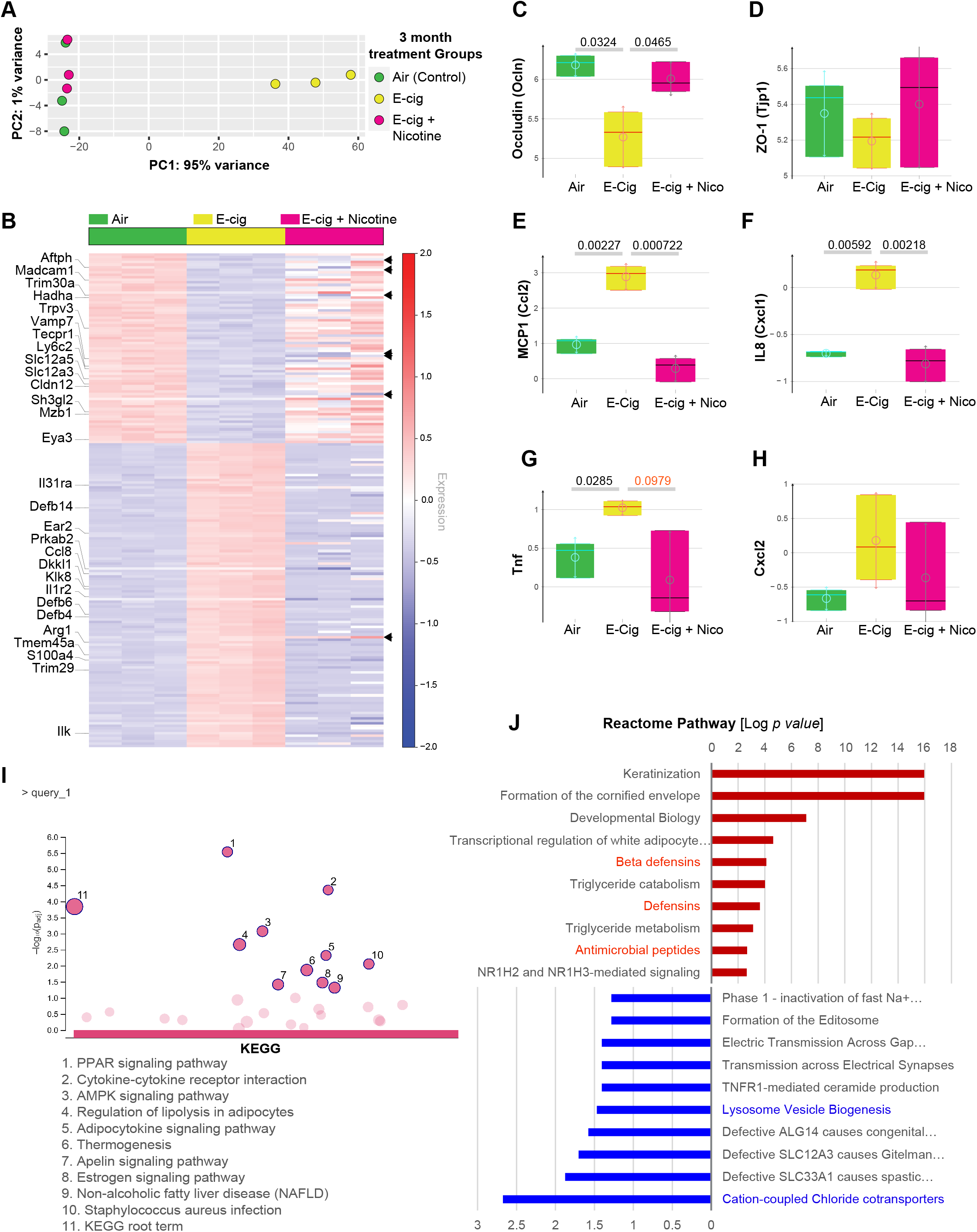
Nicotine-free, but not nicotine-containing e-cigarettes, trigger wide-ranging changes in gene expression in the colon. **A.** Principal component analysis (PCA) of the topmost variable genes in the distal colons of the 3 groups of mice, all exposed to air or e-cig as indicated for 3 mo. PCA identified air control and nicotine-containing e-cig group as similar to each other, but distinctly different from the e-cig alone group along the first principal component. **B**. Heatmap visualization of the differentially expressed genes between the 3 groups of mice. Each row represents one of the genes, while columns represent expression averages of replicates for each investigated group. Red color indicates relative over-expression, while blue color indicates relative under-expression. Top genes in the Reactome pathway analyses are marked on the left side. Arrowheads on the right side indicate a few genes that remained altered in both e-cig alone and e-cig + nicotine groups (see **Fig S2B)**. **C-H.** Whisker plots display the levels of expression of the genes, as determined by RNA seq encoding tight junction markers (occluding, C; ZO1, D) and pro-inflammatory cytokines (Mcp1, E; Il-8, F; Tnfα, G; Cxcl2, H). **I-J**. KEGG (I) and Reactome (J) pathway analyses of the list of differentially expressed genes (see **Table S1**) reveal the most up or downregulated pathways. Red color indicates upregulation, while blue color indicates the downregulation of gene expression. No significant enrichment of pathways was seen in the list of downregulated genes by KEGG analyses.

KEGG pathway (**Fig 2I**) and GO (**Fig S3**) analyses revealed that the most enriched disease-related pathways were those that are involved in cellular sensing and response to external stress and stimuli (peroxisome proliferator-activated receptors, PPAR and 5’ AMP-activated protein kinase, AMPK signaling), cell death and programmed cell death, defense response to other organisms, metabolism (lipolysis, adipocytokine, thermogenesis) and inflammation (cytokine and receptors). With regard to cytokine signaling, we noted that *il31ra* (subunit for IL6R), *il1r2* (decoy receptor for IL1R1), *ccl8* (a chemoattractant), ear2 (chemoattractant), and *ilk* (activator of NFkB) were upregulated, whereas *trim30a* (a suppressor of NFκB) and *madcam1* (a cell-adhesion molecule required for leukocyte trafficking) were downregulated. No KEGG pathways were significantly enriched among the downregulated genes. A Reactome pathway (**Fig 2J**) analysis on the same gene sets showed enrichment of anti-microbial peptides, specifically beta-defensins (*Defb4, 6 and 14*), and de-enrichment of regulators of lysosome biogenesis (*sh3gl2, eya3, vamp7*) and chloride transporters (*slc12a3, slc12a5*). Besides these statistically enriched pathways and processes, it is noteworthy that several cancer-related genes (*dkkl1, trim29, s100a4, tmem45a,* and *klk8*) were also upregulated (**Fig 2B**), and as expected, an enrichment of pathways for differentiation program in the colon (e.g., multiple Keratins; **Fig 2J**). Many of these differentiation-related genes continued to remain high despite nicotine (**Fig S2B**). These findings suggest the upregulation of two opposing programs in the colon that tightly regulate oncogenesis in the colon.

### Chronic exposure to e-cigarettes disrupts the integrity of the human gut barrier and triggers inflammation

To translate the relevance of our findings in mice colon to the human gut, we generated organoids from colonic and ileal biopsies obtained during colonoscopy from healthy human subjects (3 independent donors, age range-29 to 71 years). These organoids were subsequently differentiated into EDMs and exposed either acutely (single exposure) or chronically (repeated exposures x 3, each 4 hours apart) prior to analyzing them at 24 hours for barrier integrity and markers of inflammation using multiple modalities (see **Fig 3A**). An acute single exposure to e-cig was associated with a significant drop in TEER (−48.1±5.5%) at 4 hours (**Fig 3B**; *left*). However, much of that initial drop at 4 hours was virtually reversed after 24 hours to levels that were similar to untreated or air-treated control EDMs (UN = 41.7±8.0%, Air = 49.1±14.5% and e-cig = 23.6±18.5% (**Fig 3B**; *right*). When the EDMs were subjected to 3x e-cig exposures, which is a more physiologic exposure based on human use patterns, the drop in TEER was sustained at 24 hours (−59.96±3.82%; **Fig 3B**; *right*).

**Figure 3:**
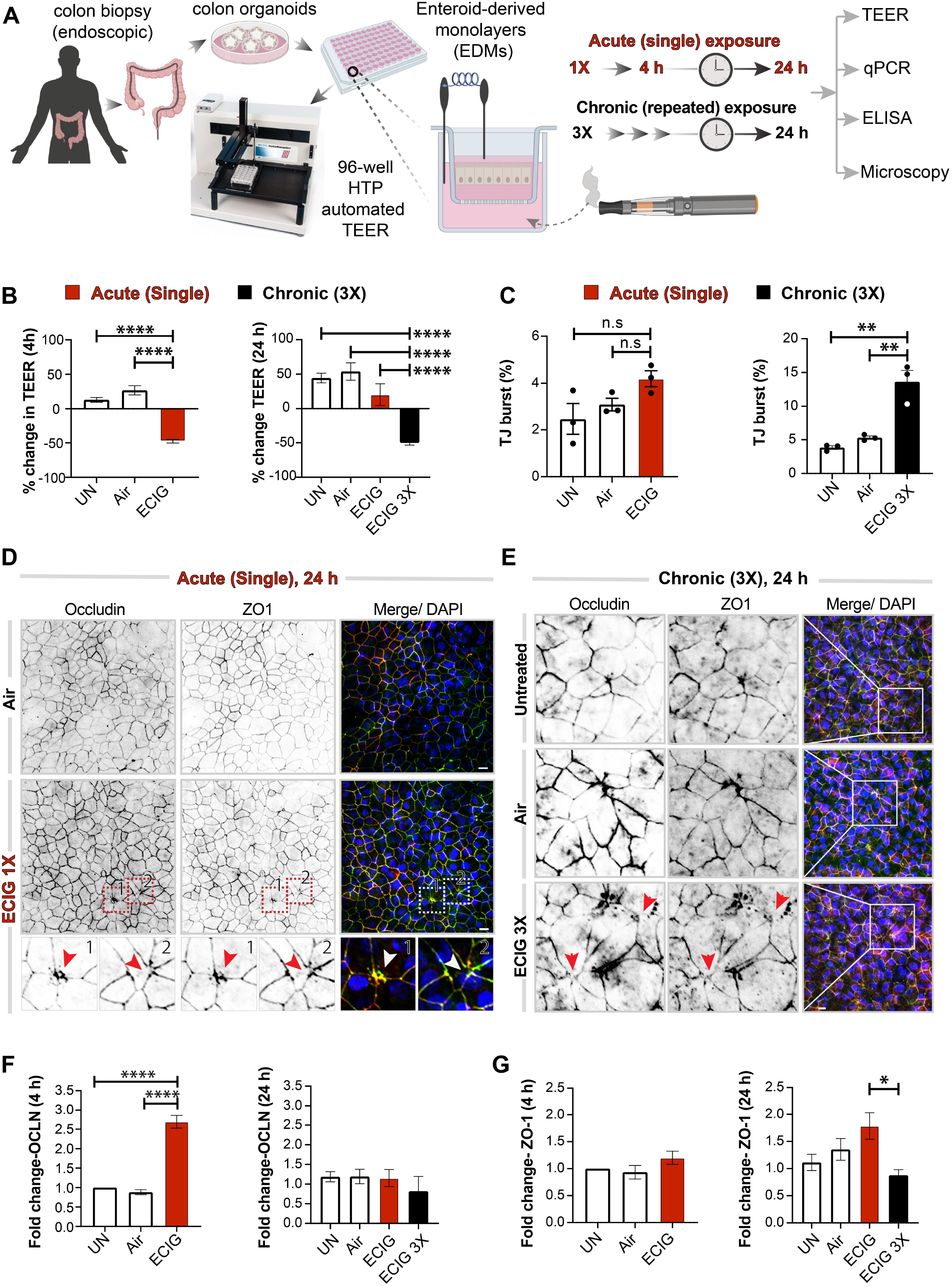
Chronic exposure to e-cigarettes disrupts the integrity of the human gut barrier, triggers inflammation. **A.** Schematic displays the key aspects of *ex vivo* disease modeling to interrogate the impact of vaping on the human colonic epithelial barrier. Two modes of exposures (acute and chronic) were modeled and a variety of functional readouts were analyzed. **B**. Bar graphs display % change in transepithelial electrical resistance (TEER) over time. Data is displayed as mean ± SEM (n = 3). Statistical significance was estimated using one-way ANOVA with Tukey’s test; ****p <◻0.0001. **C-E.** EDMs were treated as indicated prior to fixation, and then stained for tight junction (TJ) marker occludin (green), ZO1 (red) and DAPI (blue, nuclei) and analyzed by confocal microscopy. Bar graphs in **C** display the % increase in tight junction (TJ) ‘bursts’. Data is displayed as mean ± (n = 3-5 fields/condition). Statistical significance was estimated using one-way ANOVA with Tukey’s test; **p <◻0.01; n.s. = not significant. Confocal microscopic images representative of EDMs after 24 h of treatment with either a single (D) or repeated (3x; E) exposure(s) to air (control) or nicotine-free e-cig vapor-infused media are shown. Scale bar = 10 μm. **F-G**. Bar graphs display the relative fold change in mRNA expression of tight junction markers (Occludin and ZO1) in human EDMs after acute (4 h) or chronic (24 h) exposure of e-cig vapor-infused media. Data is displayed as mean ± SEM (n = 3). Statistical significance was estimated using one-way ANOVA with Tukey’s test; *p◻<◻0.05 and ****p <◻0.0001.

We also confirmed that the observed drop in TEER in the colon-derived EDMs was also associated with a loss of structural integrity of the TJs. An acute single exposure was associated with only infrequent aberrant tricellular TJ morphology and a statistically insignificant ‘burst’ appearance at 24 h (**Fig 3C**, *left*; **Fig 3D**, *arrowheads*). However, repeated 3x exposures resulted in a significant ~3-fold increase in the % of ‘burst’ TJs (**Fig 3C**, *right*; **Fig 3E**, *arrowheads*). The levels of transcripts of the membrane-integral TJ-marker, occludin, increased after acute exposure (**Fig 3F**, *left*; **Fig S4**), but returned to normal levels at 24 h (**Fig 3F**; *right*). The levels of the peripheral TJ-marker ZO1 was unchanged at 4 h after an acute exposure (**Fig 3G**; *left*), while there was a significant drop in gene expression of ZO1 after chronic repetitive multiple exposure (**Fig 3G**; *right*).

Because prior studies have implicated loss of epithelial barrier integrity as permissive to inflammation (33), we next investigated inflammatory gene expression in the e-cigarette-exposed EDMs by qPCR. Compared to untreated or air-treated controls, chronic repetitive exposure to e-cigarettes (but not single acute exposure) increased expression of all pro-inflammatory cytokines tested (**Fig 4A-D**). ELISA studies conducted on EDM supernatants confirmed that pro-inflammatory cytokine IL-8 was significantly elevated (**Fig 4E**).

**Figure 4:**
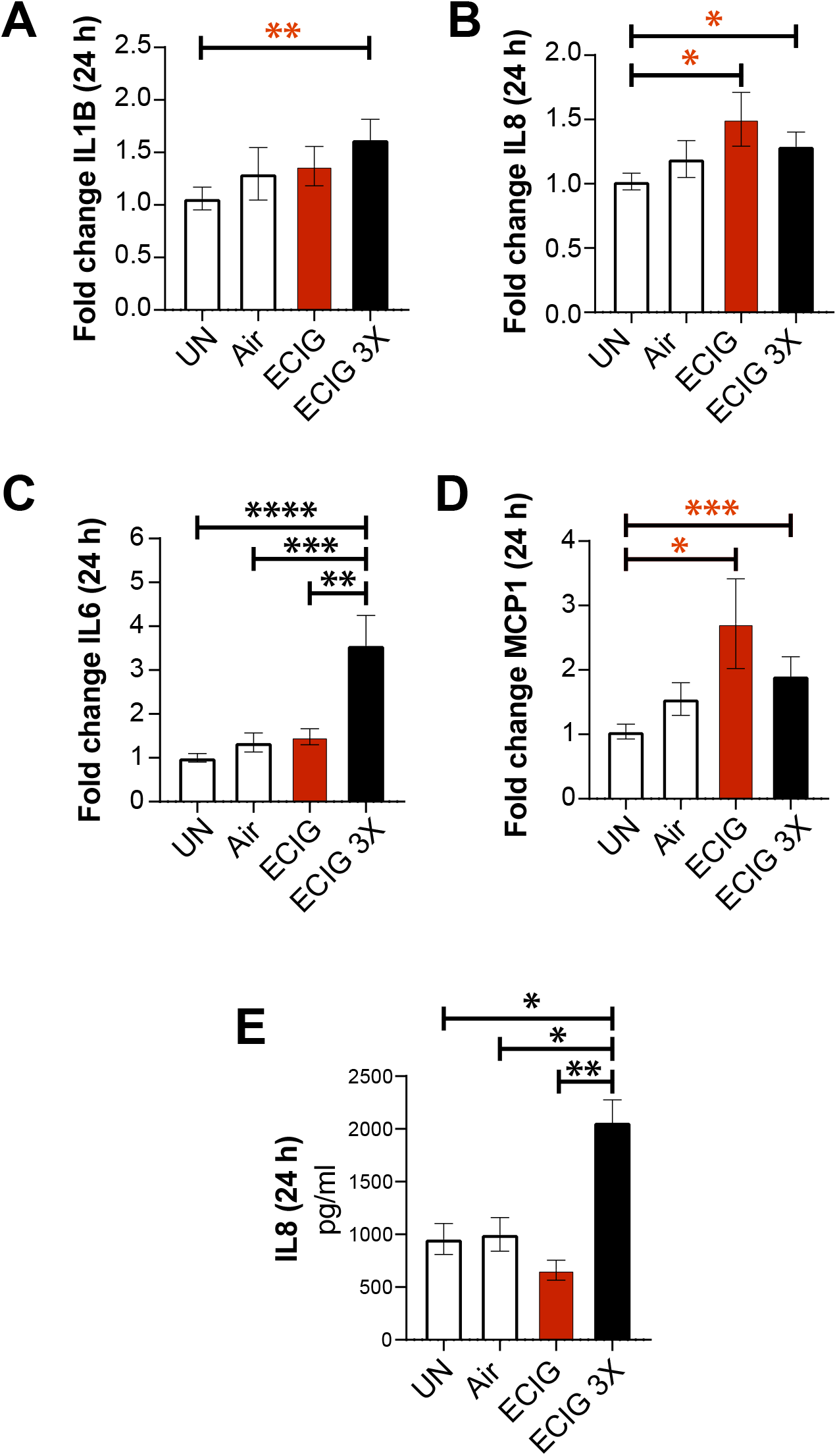
Chronic exposure to nicotine-free e-cigarette induces the expression of pro-inflammatory cytokines in the human colonic epithelium. **A-D**. Bar graphs display the relative fold change in the levels of mRNA for pro-inflammatory cytokines. Data is shown as mean ± SEM (n = 3). Statistical significance was estimated using either one-way ANOVA with Tukey’s test (black) or Mann-Whitney test (red); *p◻<◻0.05, **p <◻0.01 and ***p <◻0.001. **E**. Bar graphs display the concentration of IL-8 released in the basolateral compartment of polarized EDMs after exposure to e-cig vapor-infused media. Data is shown as mean ± SEM (n = 3 independent experiments). Statistical significance was estimated using one-way ANOVA with Tukey’s test; *p◻<◻0.05 and **p <◻0.01.

Similar findings were also observed in the case of human ileum-derived EDMs. TEER dropped significantly at 4 hours after both single and 3x exposures (**Fig S5A-C**; ~55% drop compared to control EDMs). The patterns of change in occludin and ZO1 were mirrored in the case of human ileum-derived EDMs (**Fig S5D-G**). While a single acute exposure had little or no effect on cytokine transcripts (**Fig S5H, J, L, N**), chronic repetitive e-cigarette exposure caused increased IL-1B (**Fig S5I**), IL-6 (**Fig S5K**), IL-8 (**Fig S5M**) and MCP1 (**Fig S5N**).

Taken together, these physiologic (TEER), morphologic (‘burst’ appearance of TJ’s), and transcriptomic (qPCR assessment of markers of TJ-transcripts) readouts are all in agreement, i.e., exposure to nicotine-free e-cigarette aerosols causes epithelial barrier dysfunction in the human gut. These data also demonstrate that chronic (repetitive), but not acute (single), exposure is necessary for such disruption. Furthermore, we found that disruption in barrier integrity in the setting of chronic repetitive exposure is also associated with the induction of pro-inflammatory cytokines.

### Exposure of the gut epithelium to e-cigarette aerosols accentuates inflammatory responses to infections

Because the gut epithelial barrier of those who vape is concomitantly exposed to chemical components of e-cigarettes as well as luminal microbes, we exposed EDMs simultaneously to both stressors. First we exposed the basolateral side of EDMs grown on transwells to e-cigarette-infused media (mimicking the absorption of core chemicals contained within nicotine-free vaping aerosols into the blood stream and diffusion into tissues) and subsequently challenged the apical surface with live pathogenic microbes (to simulate luminal microbes) (**Fig 5A**). We used the adherent invasive *E. coli (AIEC)*-*LF82,* a pathogenic strain that was originally isolated from a patient with IBD (34). Compared to untreated controls, EDMs repeatedly exposed to e-cigarette aerosol media had a significant drop in the levels of occludin mRNA (**Fig 5B**; *left*) and significant increases in the levels of transcripts of inflammatory cytokines IL-1B, IL-8 and MCP1 (**Fig 5C-E**). Levels of IL-6 also trended up but fell short of statistical significance (**Fig 5F**). ELISA studies confirmed that EDMs repeatedly exposed to e-cigarette aerosol media also secreted higher amounts of IL-8 (**Fig 5G**) and MCP1 (**Fig 5H**). Unlike the EDMs that were repeatedly exposed to e-cigarettes (3x), those exposed only once did not have a significant reduction in occludin (**Fig 5B**; *left*), nor induction of proinflammatory cytokines (**Fig 5C-H**). These findings indicate that exposure to the common core chemical components of e-cigarette aerosols is sufficient to make the gut hyperreactive to microbes, and that repetitive exposure is necessary for such hyperresponsiveness.

**Figure 5:**
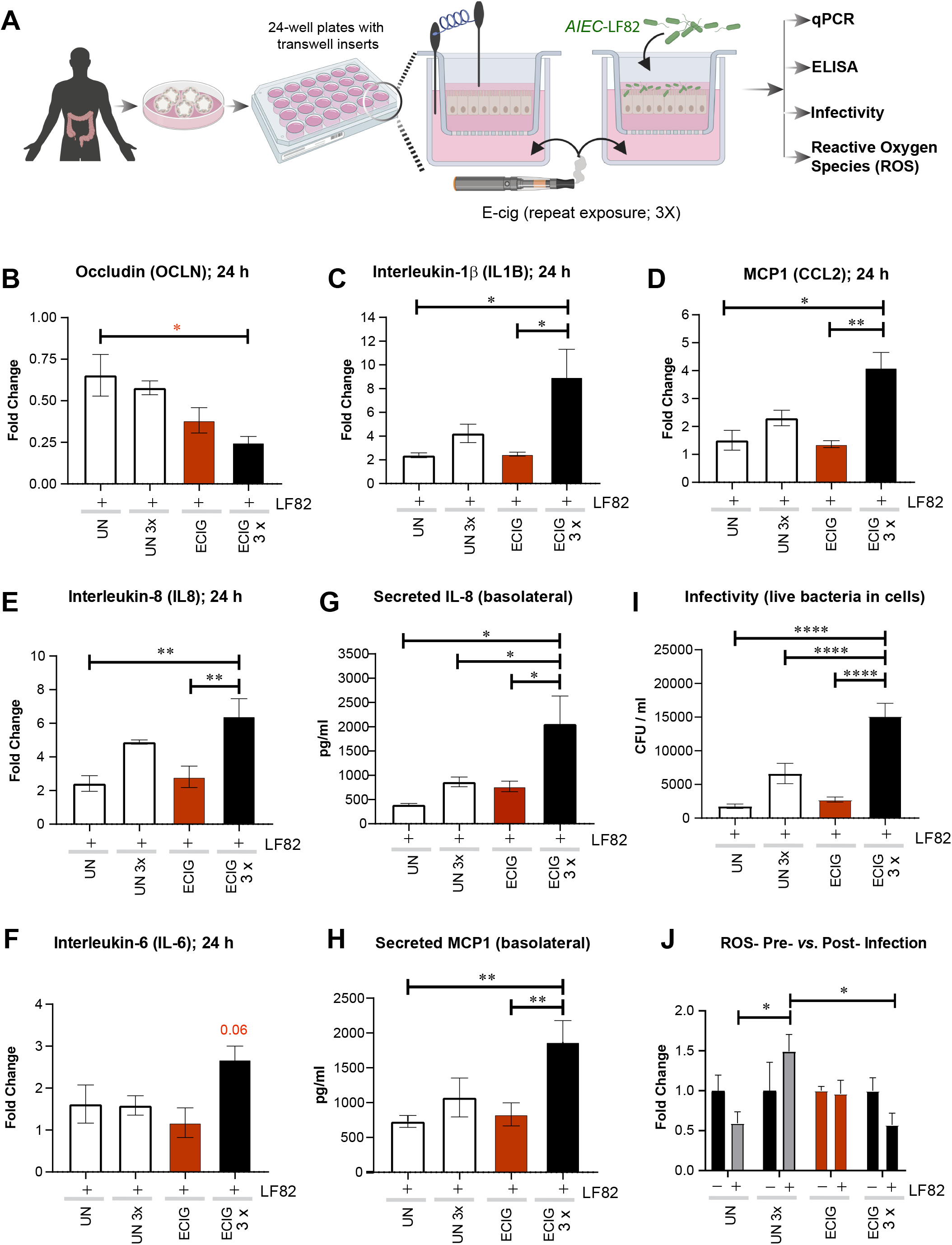
Chronic exposure to nicotine-free e-cigarette induces the expression of pro-inflammatory cytokines in the human colonic epithelium. **A.** Schematic displays the overall experimental design for assessing how e-cig affects the gut epithelial response to infections pathogenic microbes, e.g., *E. coli* strain AIEC-*LF-82*.. **B-F**. Bar graphs display the relative fold change in the levels of mRNA for pro-inflammatory cytokines. Data is shown as mean ± SEM (n = 3 independent experiments). Statistical significance was estimated using one-way ANOVA with Tukey’s test; *p◻<◻0.05, **p <◻0.01, ***p <◻0.001 and ****p <◻0.0001. **G-H.** Bar graphs display the concentration of IL-8 (G) and MCP-1 (H) released in the basolateral compartment of polarized EDMs after exposure to e-cig vapor-infused media. Data is shown as mean ± SEM (n = 3 independent experiments). Statistical significance was estimated using one-way ANOVA with Tukey’s test; *p◻<◻0.05 and **p <◻0.01. **I**. Bar graphs display the bacterial load internalized in EDMs pretreated as indicated with or without single or repeated exposure to e-cig vapor-infused media and then exposed to pathogenic *AIEC-LF82* for 3 h. Data is expressed as no. of internalized bacteria to the infected control EDMs (untreated; i.e., not exposed to e-cig) and is represented as the mean ± SEM of three separate experiments. Statistical significance was estimated using one-way ANOVA with Tukey’s test; **p◻<◻0.01. **J**. Bar graphs display cellular accumulation of ROS, as determined by measuring the levels of oxidized DNA in the supernatant in the basolateral compartment of polarized EDMs after exposure to the indicated treatments. Data is represented as the mean ± SEM of three separate experiments. Statistical significance was estimated using one-way ANOVA with Tukey’s test; *p◻<◻0.05.

Because cytokine production by epithelial cells after infection is a culmination of multiple events during epithelial sensing and signaling that are triggered by microbes, we next asked how exposure to e-cigarettes impact some of the early steps, i.e., infectivity of the gut epithelium and epithelial reaction to such infection by production of reactive oxygen species (ROS); the latter serves as a critical second messenger which modulates innate immune signaling in the gut epithelium (35). EDMs exposed repeatedly to e-cig (3x) showed statistically significant higher number of internalized bacteria compared to control EDMs after 3 hours of infection, demonstrating decreased host defenses with higher infectivity of gut epithelium after e-cigarette exposure (**Fig 5I**). Finally, we found that repeated exposures of e-cigarette aerosol media (3x) followed by infection of EDMs was associated with reduction in ROS (**Fig 5J**). Unlike the chronically exposed EDMs, those exposed only once did not show a significant increase in infectivity; nor did they show a significant reduction in ROS production (**Fig 5I-J**). These findings indicate that chronic e-cigarette use may lead to diminished host defenses in the gut, namely reduced ROS, leading to increased susceptibility to bacterial infections.

Taken together, these findings indicate that chronic repetitive exposure to e-cigarettes alter the gut epithelial cell response to infection with pathogenic microbes, characterized by higher infectivity and induction of proinflammatory cytokines and a failure to induce ROS. Because the overall composition of the gut microbes does not appear to be significantly altered among the subjects who consume e-cigarettes (26), our findings show that e-cigarettes may impair gut homeostasis primarily *via* modulation of host responses to microbes.

## DISCUSSION

### E-cigarettes trigger gut inflammation

The major discovery we report in this work is that chronic repetitive but not acute exposure to e-cigarette aerosols disrupts the gut epithelial barrier, increases the susceptibility of the gut lining to bacterial infections and triggers gut inflammation. We also pinpoint the components in the e-liquid as the major culprit. We established causality by using near physiologic *ex vivo* murine and human gut models; the minimalistic nature of the enteroid monolayer system and our ability to manipulate it sequentially and in a physiologically relevant manner (context, dimension and orientation) allowed us to pinpoint the target cell for e-cig-induced injury as the gut epithelial cell. By using invasive *E. coli* in co-culture studies with EDMs, we also determine that handling of microbes by the gut epithelium was fundamentally impaired upon chronic and repetitive exposure to e-cig, resulting in higher infectivity and inflammation. Our findings are in keeping with prior studies showing higher infectivity and inflammation in the epithelial lining of the oral mucosa (25) and the lung (36–38).

### E-cigarettes broadly impact gut health

Other major outcomes of this study are the insights gained from the extent and nature of the altered transcriptional landscape of the murine gut that is repeatedly exposed to e-cig over months. Our RNA seq studies showed three major inter-related themes of altered transcriptional programs. First, are pathways concerning cellular response to stress and stimuli; prominent inductions of genes that participate within the PPAR and AMPK signaling pathways. Second, are pathways concerning mucosal response to infection and inflammation, with prominent induction of genes that encode the anti-microbial peptide β-defensins and downregulation of genes that concern lysosomal biogenesis and genes like *Tecpr1,* which is necessary and sufficient for autophagic clearance of microbes (39). The upregulation of stress response genes and the very specific pattern of upregulation of β-isoform of defensins is not unique to the gut; transcriptomic analyses on human bronchial epithelium have documented the same previously (40). The third and final theme is that of a balanced upregulation of genes that support pro- and anti-oncogenic pathways and processes, the most prominent of which were genes involved in cellular differentiation, i.e., multiple keratins. Because, a sufficient amount of keratin is needed for efficient stress protection in the colonic epithelia and because keratins play an essential role of maintaining epithelial barrier and its down-regulation in intestinal tissue has been correlated with the progression of IBD (41, 42), our findings suggest that the three themes of altered gene expression may be interrelated consequences of epithelial stress response to chronic stimuli (simultaneous exposure to e-cig and microbes) and inflammation.

### E-liquid, not nicotine, is the culprit

Our RNAseq studies using murine models of vaping unexpectedly but decisively revealed that it is the e-liquid component in e-cigarettes that induces broad and sweeping changes on gene expression in the gut. Virtually all changes are reversed by a concomitant co-administration of nicotine. These findings are in keeping with the barrier-tightening effect and anti-inflammatory effect of nicotine that have been demonstrated previously. For example, exposure of epithelial monolayers to nicotine and its metabolites, at concentrations corresponding to those reported in the blood of smokers significantly improves tight junction integrity, and thus, decreases epithelial gut permeability (43). Similarly, studies in humans (44, 45) have shown that nicotine does tighten the gut and that such effect may only be seen upon chronic, but not acute exposures (46–48). As for mechanism(s) behind such tightening, upregulation of TJ markers (43, 49), most prominently that of occludin, has been reported. Others have shown that tightening of the gut barrier also translates to reduced inflammation in the gut mucosa; nicotine appears to have a direct anti-inflammatory effect on monocytes (50–53) and T cells (54, 55), potentially through direct activation of the cholinergic anti-inflammatory pathway, which involves inhibition of NFkB signaling (56). Furthermore, nicotine has been generally found to serve as a protective factor for the development and progression of ulcerative colitis (UC), a condition that is characterized by leakiness of the gut barrier and chronic inflammation in the gut lining. That the nicotine-containing e-cig group is protected from the impacts of e-cig is most consistent with the possibility that the addition of the barrier-protective agent, nicotine, to the e-cig formulation was sufficient to most optimally balance the barrier-destructive effects of e-cig alone (showcased here).

### Study limitations

The safety of flavors and other additives (i.e., cannabinoids, THC, etc) was not assessed here. Designed to appeal to youth, women and minorities, the flavorants make up 20-60 of the chemicals contained within e-liquids. In the absence of regulations over e-liquid contents and compositions, and the plethora of flavors there is to test (thousands), scaling up to power such a study is not trivial. Another limitation is that the enteroid system is a minimalistic model, and lacks critical components (e.g., immune and non-immune cells) that are important for setting up a vicious cycle of inflammation in the gut and further damage of the epithelial lining. Thus, it is possible that the phenotypes we observed in our EDM-based assays underestimate the full extent of injury and inflammation due to e-cigarettes. Finally, although we reconstitute the EDM model with live microbes (the single pathogenic strain of *AIEC-LF82*), the full impact of e-cigarettes in the setting of the polymicrobial gut luminal milieu was not estimated here. Ongoing work is investigating several of these outstanding aspects.

**Figure 6:**
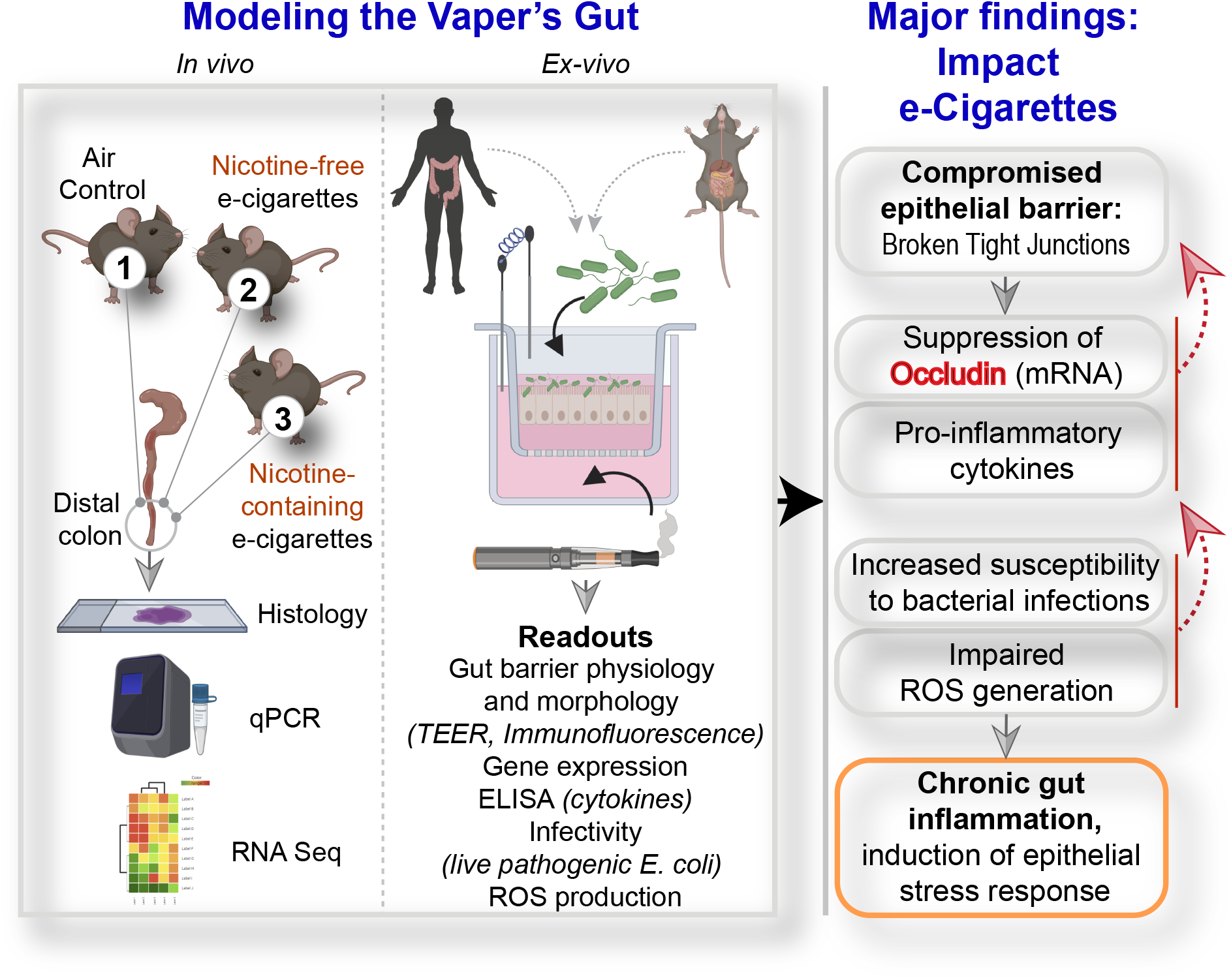
Summary of findings. *<u>Left:</u>* Schematic summarizes the various *in vivo* and *ex vivo* murine and human models used (top) and the readouts assessed (bottom) in this work to study the impact of vaping on the gut barrier. *<u>Right</u>* The major findings and conclusions from this study, with the proposed hierarchy of events over time (from top to bottom), beginning with epithelial barrier disruption, which is permissive to altered gene expression (lower occludin and higher cytokines) and heightened susceptibility for and response to bacterial infections, culminating in chronic inflammation and epithelial stress response. Potential feed-forward loops are indicated as red interrupted arrows.

## MATERIALS & METHODS

All methods involving human and animal subjects were performed in accordance with the relevant guidelines and regulations of the University of California San Diego and the NIH research guidelines. H&E staining, Human Subjects, Isolation of organoids, Preparation of EDMs, RNA isolation, qPCR and RNA Sequencing, Immunofluorescence, assessment of oxidative DNA/RNA damage are detailed in *SI Appendix*, and briefly mentioned here.

### Murine E-Cigarette aerosol exposures

Six to eight-week old male or female C57BL/6 (Envigo) were acclimatized to the SciReq whole body exposure inhalation system for 30 min a day for 3 days prior to beginning e-cigarette exposures. Mice were placed into individual slots in the whole-body exposure system and exposed for 4 sec every 20 sec for 1 h/day, 5 d/wk, for 1 wk or 12 wks. Mice received e-cigarette vapor produced from e-liquid containing 70/30 propylene glycol and glycerol with nicotine (Sigma) at a concentration of 6 mg/mL or without nicotine. At the end of the exposure period, mice were anesthetized with 10mg/kg and 100 mg/kg of xylazine and ketamine, respectively.

### Preparation of E-cigarette vapor-infused media

E-liquid mixture of 70% propylene glycol and 30% glycerol and 6mg/mL nicotine (purchased from Sigma) without flavors or additives were used. E-cigarette atomizer and battery were obtained from Scireq containing a Kangertech Subtank Plus (7mL), using a 0.15 ohm coil, and rechargeable battery. Fresh e-cig vapor-infused media was created by activating the battery via application of negative pressure by the InExpose system (SciReq), the e-liquid was heated and drawn through the internal atomizer and then into a 60 ml syringe containing 10 ml of wash media (DMEM/F12 with HEPES, 10% FBS). The wash media was exposed to 50mLs of e-cig vapor generated from the vaporization of the e-cig liquid, (with or without 6mg/ml nicotine) followed by a 12-second shake; this is repeated 30 times.

### Human Subjects

To generate healthy enteroids, a fresh biopsy was collected from healthy subjects undergoing routine colonoscopy using the protocol approved by the Human Research Protection Program Institutional Review Board (Project ID# 190105).

### Isolation of organoids and preparation of Enteroid-derived monolayers (EDMs)

Intestinal crypts, comprised of crypt-base columnar (CBC) cells, were isolated from both colonic and ileal tissue specimens using the previously published paper (33, 57). EDMs were generated on either 24-well or 96-well transwells with a 0.4 μm pore polyester membrane (Corning) as described previously (33, 57, 58).

### The treatment of enteroid-derived monolayers (EDMs) with E-cigarette for functional assays

The polarized differentiated EDMs were treated with e-cigarette vapor infused media, added to the basolateral compartment of the transwells for the indicated times. The EDMs were used to measure transepithelial electrical resistance (TEER), stained for immunofluorescence or the preparation of RNA and for the collection of supernatants from apical and basolateral sides.

### The measurement of Transepithelial electrical resistance (TEER)

Two different methods were used for the measurement of TEER in manual low-(LTP) and automated high-throughput (HTP) modes. For manual assessments, TEER was measured in 24-well transwell plates and for automated TEER, 96 well plates were used to measure the values at 0 h, 1 h, 4 h, 8 h, and 24 h, following exposure to e-cig vapor infused media using the STX2 electrodes with digital readout by EVOM2 (WPI). For automated assessments, TEER was measured in 96-well transwell plates with EDMs was measured using WPI automated TEER Measurement System (REMS AutoSampler, Version 6.02).

### Co-cultures of EDMs with invasive *E. coli*

Adherent Invasive *Escherichia coli* strain LF82 (*AIEC-* LF82), isolated from the specimens of Crohn’s disease patient, was obtained from Arlette Darfeuille-Michaud (59). Co-cultures were carried out as done previously and expanded in SI(33, 57).

### Measurement of DNA/RNA oxidative damage

The level of DNA damage in EDMs following exposure to air and e-cigarette vapor infused media was quantified with the DNA/RNA Oxidative Damage ELISA Kit (Cayman Chemical, USA) according to the manufacturer’s instruction and previous papers (60, 61).

### Immunofluorescence staining

After the final TEER measurement of all EDM treatments, the media was removed from apical and basolateral compartments. The exposed EDMs were fixed, permeabilized and stained for occluding and ZO-1 and imaged using a Leica TCS SP5 Confocal Microscope as done previously(33). Z-slices of a Z-stack were overlaid to create maximum intensity projection images; all images were processed using FIJI (Image J) software.

### Statistical analysis

TEER, qPCR and ELISA results were expressed as the mean ± SEM and compared using a one-way ANOVA with Tukey’s test or Mann-Whitney test. Results were analyzed in the GraphPad Prism and considered significant if p-values were < 0.05.

## ACKNOWLEDGMENTS

We thank the UC San Diego HUMANOID Center of Research Excellence (CoRE) for access to human the biobanked organoids, media and technical support. This publication includes data generated at the UC San Diego IGM Genomics Center utilizing an Illumina NovaSeq 6000 that was purchased with funding from a National Institutes of Health SIG grant (#S10 OD026929). We acknowledge the P30 grant (NIH/NIDDK, P30DK120515) that subsidized the RNA Seq and histology work showcased here. This work was supported by awards from the Tobacco-Related Disease Research Program (TRDRP): 28IP-0024 (to P.G. and S.D.), T30IP0965 (to L.C.A.) and 26IP-0040 (to L.C.A.), the National Institutes for Health (NIH) grants DK107585 (to S.D.), AI141630 (to P.G), R01HL147326 (to L.C.A.) and R00-CA151673 (to D.S). S.D. was also supported by a DiaComp Pilot and Feasibility award (Augusta University). P.G., S.D. and D.S were supported by NIH/National Center for Advancing Translational Sciences (NCATS) award UG3TR002968 and the Leonna M Helmsley Charitable Trust. S.R.I was supported by the NIH Diversity Supplement award. J. E. was supported by a Postdoctoral Fellowship from the American Cancer Society, United States, (PF-18-101-01-CSM). L.C.A. was also supported by an American Heart Association beginning grant-in-aid 16BGIA27790079, an ATS Foundation Award for Outstanding Early Career Investigators, and had salary support in part from the Veterans Affairs San Diego Healthcare System.

## AUTHOR CONTRIBUTIONS

A.S, J.L, AG.F, A.M, T.K, I.M.S, S.R.I have designed and performed the experiments; R.F.P and D.S. have performed the RNA seq analysis; J.E has performed and supervised the confocal microscopy experiments and provided scientific input. A.S, J.L, AG.F, P.G, S.D analyzed the data and generated the first draft of the manuscript; P.G and S.D wrote the manuscript. L.C.A. has provided expertise and access to the e-cigarette-infused media and e-cig treated murine colonic specimens; P.G, L.C.A and S.D supervised the project.

