## Supplementary Online Materials for "E-cigarettes compromise the gut barrier and trigger gut inflammation"

#### Supplementary Online Information

### MATERIALS AND METHODS

#### Key Resource Table

| REAGENT or RESOURCE | SOURCE | IDENTIFIER |
|---|---|---|
| **Biological Samples** | | |
| C57B/L6 mice | ENVIGO/HARLAN | |
| Adherent Invasive Escherichia coli strain LF82 (*AIEC*-LF82) | Arlette Darfeuille-Michaud, Inserm | (1) (2)<br>Darfeuille-Michaud, A |
| Human tissue biopsies | HUMANOID CoRE<br>VA hospital, San Diego | UCSD HRPP Project ID190105 |
| **Primers (Mouse/Human)** | Forward primer (3'- 5') | Reverse primer (3'- 5') |
| Mouse 18S qPCR primers | GTAACCCGTTGAACCCCATT | CCATCCAATCGGTAGTAGCG |
| Mouse ZO-1 qPCR primers | GGGAGGGTCAAATGAAGACA | GGCATTCCTGCTGGTTACAT |
| Mouse IL-6 qPCR primers | CCCCAATTTCCAATGCTCTCC | CGCACTAGGTTTGCCGAGTA |
| Mouse IL-1b qPCR primers | GAAATGCCACCTTTTGACAGT | CTGGATGCTCTCATCAGGACA |
| Mouse TNF-alpha qPCR primers | CCACCACGCTCTTCTGTCTA | AGGGTCTGGGCCATAGAACT |
| Mouse IL-8 qPCR primers | CCTGCTCTGTCACCGATG | CAGGGCAAAGAACAGGTCAG |
| Mouse MCP-1 qPCR primers | AAGTGCAGAGAGCCAGACG | TCAGTGAGAGTTGGCTGGTG |
| Mouse Occludin qPCR primers | CCTCCAATGGCAAAGTGAAT | CTCCCCACCTGTCGTGTAGT |
| Mouse Claudin-1 qPCR primers | CCCCCATCAATGCCAGGTATG | AGAGGTTGTTTTCCGGGGAC |
| Mouse Claudin-2 qPCR primers | CCTTCGGGACTTCTACTCGC | TCACACATACCCAGTCAGGC |
| Human IL-8 qPCR primers | GAGCACTCCATATGGCACAAA | ATGGTTCCTTCCGGTGGT |
| Human IL-6 qPCR primers | CCAGAGCTGTGCAGATGAGT | CTGCAGCCACTGGTTCTGT |
| Human IL-1b qPCR primers | CCACAGACCTTCCAGGAGAATG | GTGCAGTTCAGTGATCGTACAGG |
| Human MCP-1 qPCR primers | AGTCTCTGCCGCCCTTCT | GTGACTGGGGCATTGATTG |
| Human ZO-1 qPCR primers | CGGTCCTCTGAGCCTGTAAG | GGATCTACATGCGACGACAA |
| Human Occludin qPCR primers | TCAGGGAATATCCACCTATCACTTCAG | CATCAGCAGCAGCCATGTACTCT TCAC |
| **Software programs** | | |
| Prism | Graphpad | https://www.graphpad.com/scientific-software/prism/ |
| Illustrator | Adobe | https://www.adobe.com/products/illustrator.html |
| Leica Application Suite X (LAS X) | Leica Microsystems | https://www.leica-microsystems.com/products/microscope-software/p/leica-las-x-ls/ |
| Image J | FIJI | https://imagej.net/Welcome |
| AxioImager Z1 microscope | Carl Zeiss MicroImaging | |
| **Chemicals and Reagents** | | |
| Direct-zol RNA Miniprep Kit | Zymo Research | R2052 |
| Human IL-8/CXCL8 DuoSet ELISA | R&D systems | DY208 |
| Human CCL2/MCP-1 DuoSet ELISA | R&D systems | DY279 |
| Zinc Formalin Fixative | Fischer Scientific | 23313096 |
| DMEM with 10% FBS | Thermo Fisher Scientific | 11995073 |
| Fetal bovine serum | SIGMA-Aldrich | F2442-500ML |
| SB431542 (an inhibitor for TGF-β type I receptor | Bio-Techne Sales corp | 1614/50 |
| Y27632 (ROCK inhibitor) | Tocris | 1254 |
| Advanced DMEM/F12 | Thermo Fisher Scientific | 12634028 |
| Trypsin | Thermo Fisher Scientific | 15090046 |

| | | |
|---|---|---|
| Prolong Gold | Thermo Fisher Scientific | P36930 |
| DNA/RNA Oxidative Damage ELISA Kit | Cayman Chemical, USA | 501130 |
| Collagenase type II | Invitrogen | 17101015 |
| 70-µm cell strainer | Thermo Fisher Scientific | 22-363-548 |
| Matrigel | CORNING | 354234 |
| Gentamicin | Thermo Fisher Scientific | 15750060 |
| Transwell inserts | Thermo Fisher Scientific | 07-200-154 |
| qScript™ cDNA SuperMix | Quantabio | 101414-108 |
| Methanol | ACS | EM-MX0485-3 |
| Triton X-100 | SIGMA-Aldrich | X100-500ML |
| Trypan Blue | Thermo Fisher Scientific | 15250061 |
| HTS Transwell®-96 Permeable Support with 0.4 µm PET Membrane | Corning | 7369 |
| **Antibodies** | | |
| Anti-Occludin mouse monoclonal antibody (1:500) | Invitrogen | 331500 |
| Anti-ZO-1 rabbit polyclonal antibody (1:500) | GeneTex | GTX108627 |
| Alexa Fluor 594 conjugated goat anti-rabbit IgG (1:500) | Invitrogen | A11012 |
| Alexa Fluor 488 conjugated goat anti-mouse IgG (1:500) | Invitrogen | A11001 |
| DAPI (1:1000) | Invitrogen | D1306 |
| **Instruments** | | |
| Countess II Automated Cell Counter | Thermo Fisher Scientific | AMQAX1000 |
| Epithelial Volt/Ohm (TEER) Meter | WPI | SKU EVOM2 |
| Automated TEER Measurement System (REMS AutoSampler, Version 6.02) | WPI | SYS-REMS |
| Corning HTS Transwell-96 Electrode | WPI | REMS-96C |
| Leica Automated Inverted Microscope | Leica Microsystems | Leica DMI4000 B |

**RNA isolation and qRT-PCR:** RNA was isolated from mouse colon tissues and EDM samples according to manufacturer's instructions, followed by cDNA synthesis. Quantitative Real-Time PCR was conducted for target genes and normalized to housekeeping gene 18S rRNA. Fold change of treatment wells was determined relative to the control condition. Primer sequences are provided in key resource Table above.

**RNA Seq and pathway enrichment analyses:** RNASeq data was processed via kallisto (5) using the human genome build GRCh38 ensembl version 94 and corresponding genome annotation file to compute TPM (Transcripts Per Millions) (6, 7) values. We used log2(TPM+1) to compute the final log-reduced expression values. StepMiner (8) algorithm was used to compute a threshold for the high and low values for each gene. To compute the heatmap, the gene expression values were further normalized according to a modified Z-score approach centered around StepMiner threshold (formula = (expr -SThr)/3*stddev). All statistical tests were performed using R version 3.2.3 (2015-12-10). Standard t-tests were performed using python scipy.stats.ttest_ind package (version 0.19.0) with Welch's Two Sample t-test (unpaired, unequal variance (equal_var=False), and unequal sample size) parameters. Multiple hypothesis correction were performed by adjusting *p* values with statsmodels.stats.multitest.multipletests (fdr_bh: Benjamini/Hochberg principles). The results were independently validated with R statistical software (R version 3.6.1; 2019-07-05).

Over-representation (KEGG, GO, etc) analyses were carried out using the **g:GOSt** platform on p:Profiler (https://biit.cs.ut.ee/gprofiler/gost). **g:GOSt** performs functional over-representation analysis (ORA) on input gene lists and is used to detect and visualize statistically significantly enriched terms. G:GOSt curates data from Ensembl database and fungi, plants or metazoa specific versions of Ensembl Genomes, and parasite specific data from WormBase ParaSite. In addition to Gene Ontology, they include pathways from KEGG, Reactome and WikiPathways; tissue specificity from Human Protein Atlas; protein complexes from CORUM and human disease phenotypes from Human Phenotype Ontology. g:GOSt supports close to 500 organisms and accepts hundreds of identifier types.

**H& E staining:** Colonic specimens were rinsed with phosphate-buffered saline (PBS) and fixed in zinc formalin for 24 h and embedded in paraffin. Paraffin sections (4 µm) were cut and de-waxed prior to immune-histochemical staining. Sections were stained with hematoxylin/eosin (H&E). Tissue sections were evaluated and images were taken by standard light microscopy.

**Human subjects:** Human ileum and colonic biopsies were collected from healthy subjects undergoing routine colonoscopy for colon cancer screening using the protocol approved by the Human Research Protection Program Institutional Review Board (Project ID# 190105). For all the deidentified human subjects, information including age, ethnicity, gender, previous history of the disease, and the medication was collected from the chart following the rules of HIPAA. Each human participant was recruited to the study following an approved human research protocol and signed a consent form approved by the Human Research Protection Program at the University of California, San Diego. Each donor agrees that their gastro-intestinal specimens will be used to generate an enteroid line at UC San Diego's HUMANOID™ Center of Research Excellence (CoRE) for functional studies.

**Isolation of enteroids from mouse and ileum and colonic specimens of healthy human:** Intestinal crypts, comprised of crypt-base columnar (CBC) cells, were isolated from both colonic tissue of mice; and human colonic and ileal tissue specimens using the previously published paper (9, 10). Briefly, intestinal crypts were dissociated from tissues by mincing with collagenase type I (2 mg/mL solution containing gentamicin 50 ug/mL). The plate was incubated in a $CO_2$ incubator at 37°C, mixing every 10 minutes with vigorous pipetting in-between incubations, while monitoring the release of single epithelial units from tissue structures by light microscopy. Collagenase was inactivated by the addition of wash media (DMEM/F12 with HEPES, 10% FBS) and cells filtered through a 70 μM cell strainer. The filtered cell suspension was centrifuged at 200g for 5 min and the supernatant was aspirated, leaving a cell pellet. The number of viable intestinal stem cells was determined by the Trypan Blue Exclusion method using Countess II Automated Cell Counter. Epithelial units were resuspended in Matrigel and 15μl of cell-Matrigel suspension was added to the wells of a 24-well plate on ice and incubated upside-down in a 37°C $CO_2$ incubator for 10 min to allow for polymerization of the matrigel. After 10 min of incubation, 500μL of 50% Conditioned Media (CM, prepared from L-WRN cells with Wnt3a, R-spondin and Noggin, ATCC® CRL-3276™ (11) containing 10 μM Y27632 and 10 μM SB431542 were added to the suspension. For the human colonic specimens, a proprietary cocktail was added to the above media. The medium was changed every 2 days and the enteroids were expanded and frozen in liquid nitrogen for biobanking.

**Preparation of Enteroid-derived monolayers (EDMs):** To prepare EDMs, single cells from enteroids in 5% conditioned media was added to diluted Matrigel (1:40) and plated either in 24-well or 96-well transwell with a 0.4 μm pore polyester membrane as done before (9, 10). The single-cell suspension was seeded at a density of approximately $2x10^5$ cells/well (in case of 24-well) or $8x10^4$ cells/well (in case of 96-well) and EDMs were differentiated for 2-3 days in 5% CM. The media was changed every 24 hours and monitored by a light microscope to evaluate the quality of EDM. As expected, the expression of EDMs showed a significant reduction of the stemness marker lgr5 in EDMs (9, 10, 12).

**The treatment of enteroid-derived monolayers (EDMs) with e-cigarette vapor-infused media for functional assays:** The polarized differentiated EDMs were treated with e-cigarette vapor infused media, added to the basolateral compartment of the transwells for the indicated times. The EDMs were used to measure transepithelial electrical resistance (TEER), stained for immunofluorescence or for the preparation of RNA and for collection of supernatants from apical and basolateral sides.

**The measurement of Transepithelial electrical resistance (TEER):** Two different methods were used for measurement of TEER in low- (LTP) and high-throughput (HTP) modes.

*Manual, LTP*: In 24-well, TEER was measured at 0 h, 1 h, 4 h, 8 h, and 24 h, following exposure to e-cigarette vapor infused media using the STX2 electrodes with digital readout by EVOM2 (WPI).

*Automated, HTP*: The TEER of 96-well transwell plate with EDMs was measured using WPI automated TEER Measurement System. WPI REMS-96C recording electrode was used to record TEER, compatible with Corning 96-well plate format. REMS-96C recording electrode was sterilized in 70% Ethanol, followed by rinse in PBS and media. The REMS-96C apical electrode was calibrated to measure TEER approximately 1 mm above transwell membrane. Transwell-read time set to 12sec/well. Once set-up complete, plate removed from incubator and TEER measured directly afterward; the same read sequence was repeated every subsequent read to mitigate TEER artifacts due to temperature fluctuations. TEER recorded by REMS AutoSampler were saved as .txt files; raw TEER values (in Ωs), are converted to normalized TEER values by Raw TEER in ohms (Ω) x surface area of transwell in $cm^2$ = ohms. $cm^2$ (SA=0.143 $cm^2$ for 96-well and 0.33 $cm^2$ for 24-well).

**Co-cultures of EDMs with Adherent Invasive *E. coli (AIEC*-LF82*):*** Adherent Invasive *Escherichia coli* strain LF82 (*AIEC*-LF82), isolated from the specimens of Crohn's disease patient, was obtained from Arlette Darfeuille-Michaud (Darfeuille-Michaud et al., 2004). A single colony was inoculated into LB broth and grown for 8 h under aerobic conditions and then under oxygen-limiting conditions. 24h following treatment with e-cig vaped media, cells were apically infected with a multiplicity of infection (moi) of 30 for 3h. For gentamicin protection assay, bacteria were removed after 3h and treated with 200ug/ml of gentamicin for 90 minutes, followed by serial dilution in 1xPBS and plating on LB agar.

**ELISA:** Supernatant collected from basolateral sides of EDMs were tested for cytokines, including IL-8 and MCP-1. These studies were done exactly as described previously (13, 14) using biolegend kit as per manufacturer's protocol.

**Measurement of oxidative DNA/RNA damage:** The amount of oxidative DNA damage in untreated and e-cigarette treated EDMs before and after infection were quantified according to the manufacturer's instruction and previous published papers from others (15, 16) and from our group (13). In brief, this assay detects oxidized guanine species: 8-hydroxy-2′-deoxaguanosine from DNA, and 8-hydroxyguanine from either DNA or RNA.

**Immunofluorescence staining:** After the final TEER measurement of all EDM treatments, the media was removed from apical and basolateral compartments. The exposed EDMs were gently washed 3 times with room temperature PBS, fixed with cold 100% methanol at 20°C for 20 min. Methanol was removed and washed with blocking buffer (0.1% Triton TX-100, 2 mg/mL BSA diluted in PBS). The permeabilized EDM was incubated with the following primary antibodies: ZO-1 (1:500) and Occludin (1:500). Primary antibodies were removed and washed with PBS 3 times for 5 min each time; after which, the following secondary antibodies were added: Alexa Fluor 594 conjugated goat anti-rabbit IgG, Alexa Fluor 488 conjugated goat anti-mouse IgG and DAPI. Secondary antibodies were removed and washed with PBS 3 times for 5 min each time. Prolong Gold antifade reagent was added, and monolayers were stored at 4°C until imaged. Confocal Microscope with a 40x objective lens was used to image the IF stained EDMs. Z-stack images were acquired by successive 1 μm depth Z-slices of EDMs in the desired confocal channels. Fields of view that were representative of a given transwell were determined by randomly imaging 3 different fields. Z-slices of a Z-stack were overlaid to create maximum intensity projection images; all images were processed using FIJI (Image J) software.

### SUPPLEMENTARY TABLES

#### **List of upregulated genes** (n = 75)

| | | | | | | | | | |
|---|---|---|---|---|---|---|---|---|---|
| B830042I05RIK | OTUD4 | D430018E03RIK | GM20476 | PKN2 | AFTPH | SCYL3 | EHF | 1700028B04RIK | GM25219 |
| MADCAM1 | GM43213 | GM17733 | 4732463B04RIK | ASXL2 | IGHV 1-26 | TRIM30A | HADHA | AW822252 | SMG1 |
| GM31774 | ZBED6 | CDK13 | PTAR1 | TRIM36 | PRPF40A | SLC33A1 | ACNAT1 | ZFP871 | TRPV3 |
| GM12511 | RPS10-PS1 | HAO2 | 1700120E14RIK | RDH16F2 | NAIP3 | RANBP2 | ZAN | ABCG2 | TNPO1 |
| GM26812 | VAMP7 | GM11945 | MAP3K2 | TECPR1 | LY6C2 | TMEM181B-PS | ETFDH | GM26711 | SLC12A5 |
| ZFP748 | SLC12A3 | GM44216 | AL611930.1 | ALG13 | NUFIP2 | GP2 | SIRT3 | PREPL | GM15693 |
| CLDN12 | RBM45 | SH3GL2 | MZB1 | USP53 | RCOR1 | UNC13B | ATP6V0A2 | TRIML1 | GM9938 |
| MOB1B | MYB | A1CF | EYA3 | GAL3ST2c | | | | | |

#### **List of downregulated genes** (n = 120)

| | | | | | | | | | |
|---|---|---|---|---|---|---|---|---|---|
| CNIH2 | SBSN | WFDC12 | GM15519 | TMPRSS11A | STFA1 | CTCFLOS | TNNC2 | ADIG | CYP1A1 |
| LCE1D | SERPINB12 | GM26751 | PIN1 | FAM178B | ENDOU | IL31RA | 1700003F12RIK | CCDC27 | DAPL1 |
| MCPT4 | GM32772 | CNFN | KRT13 | KRT10 | DEFB14 | CSTA1 | MARC1 | GM15551 | GM94 |
| GM12496 | RETN | GP1bb | KRT6B | LGALS7 | EAR2 | BORCS5 | Hp | GM10108 | LCE1A2 |
| LEP | CRCT1 | GM48007 | 9330161L09RIK | AC164431.1 | HOXD3OS1 | LCE3F | GM11772 | CALM4 | PRKAB2 |
| LCE3E | CCL8 | APOC1 | SLS36A2 | LCE3B | LCE3A | DKKL1 | KLK8 | SPRR1A | LCE3C |
| IL1R2 | TAGLN | SULT5A1 | 2300002M23RIK | LIPE | ARXES1 | SLURP1 | DEFB6 | DEFB4 | NNAT |
| SMOC1 | EYA1 | SCD1 | RTL5 | LOR | MT4 | DISC1 | NNMT | ARG1 | GM47507 |
| KRTAP20-2 | ATP2A1 | GAL3ST4 | TMEM45A | TRIM29 | TLCD2 | FLG | SERPINB1C | S100A4 | EMB |
| CAR3 | SPRR4 | SPRR3 | LCE1A1 | FABP4 | FABP5 | RNASE2B | GLB1L2 | LCE1G | FCOR |
| STFA3 | KRT1 | A230028O05RIK | KRT5 | KRT4 | FAM25C | APOL6 | KRT9 | APOBR | RNF165 |
| GM49339 | ADIPOQ | PLIN1 | ASPRV1 | ILK | ADRB3 | GM45716 | SERPINB3a | GLRX5 | BC100530 |

**Supplementary Table S1:** Differentially regulated gene clusters in mice colon vaped with e-cigarette with and without nicotine.

#### SUPPLEMENTARY FIGURES AND LEGENDS

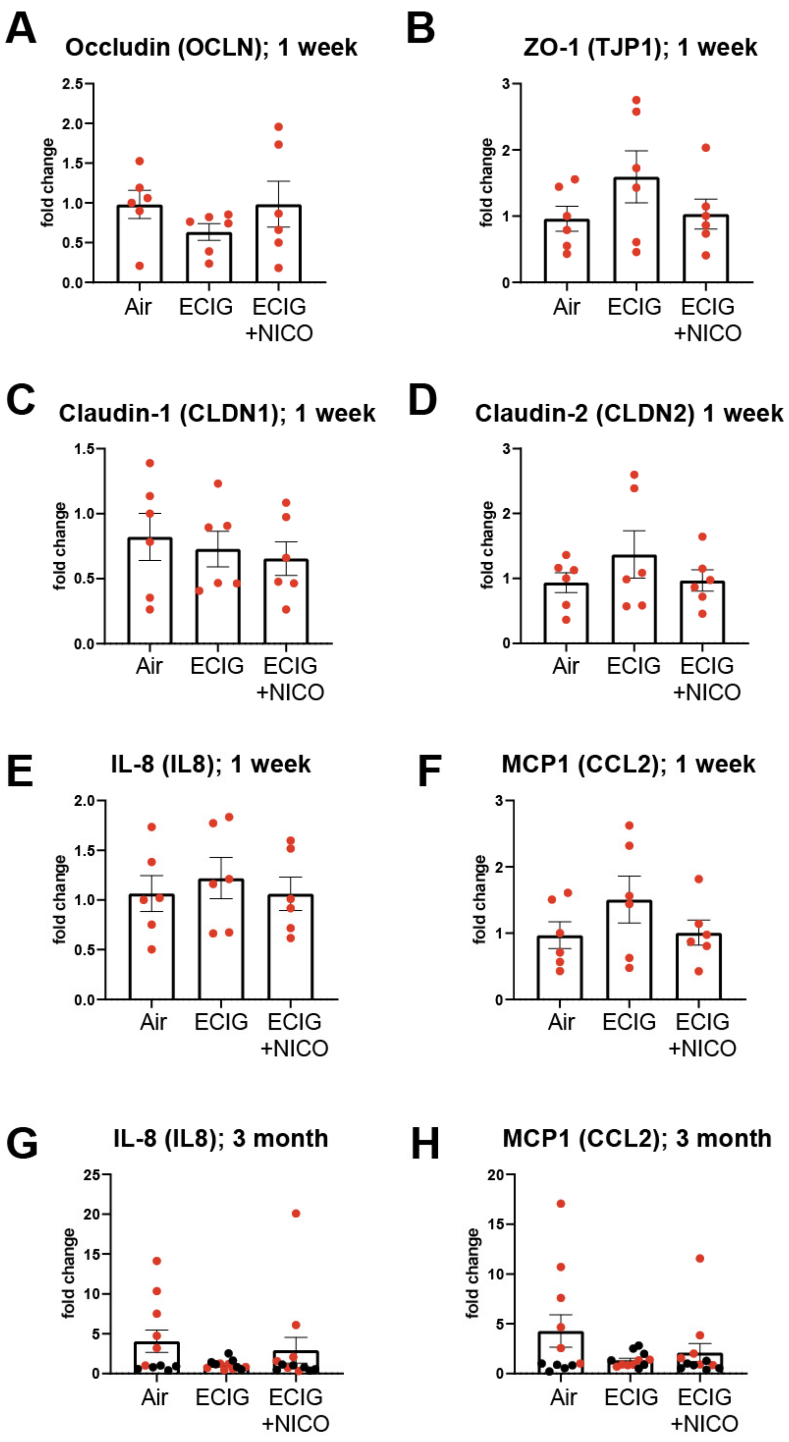

**Figure S1. Gene expression changes after acute (1 week) or chronic (3 mo) of vaping.** C57BL/6 mice were vaped using a special vaping chamber with MOD brand e-cigarette vapors (see **Fig 1A**), with (ECIG + NICO) and without (ECIG) 6 mg/ml nicotine for 1 week or 3 months by following specific regimen (details in methods), followed by isolation of the distal colon. The graphs represent the relative fold change in mRNA expression of tight junction markers (**A-D**) and pro-inflammatory cytokines (**E-H**) from 2-3 independent experiments with at least 4-5 mice/group of each experiment (Male mice = black data points; Female mice = red data points). Air = air control. Data represent as mean ± SEM. One-way ANOVA with Tukey's test was performed *$p < 0.05$, **$p < 0.01$ and ***$p < 0.001$.

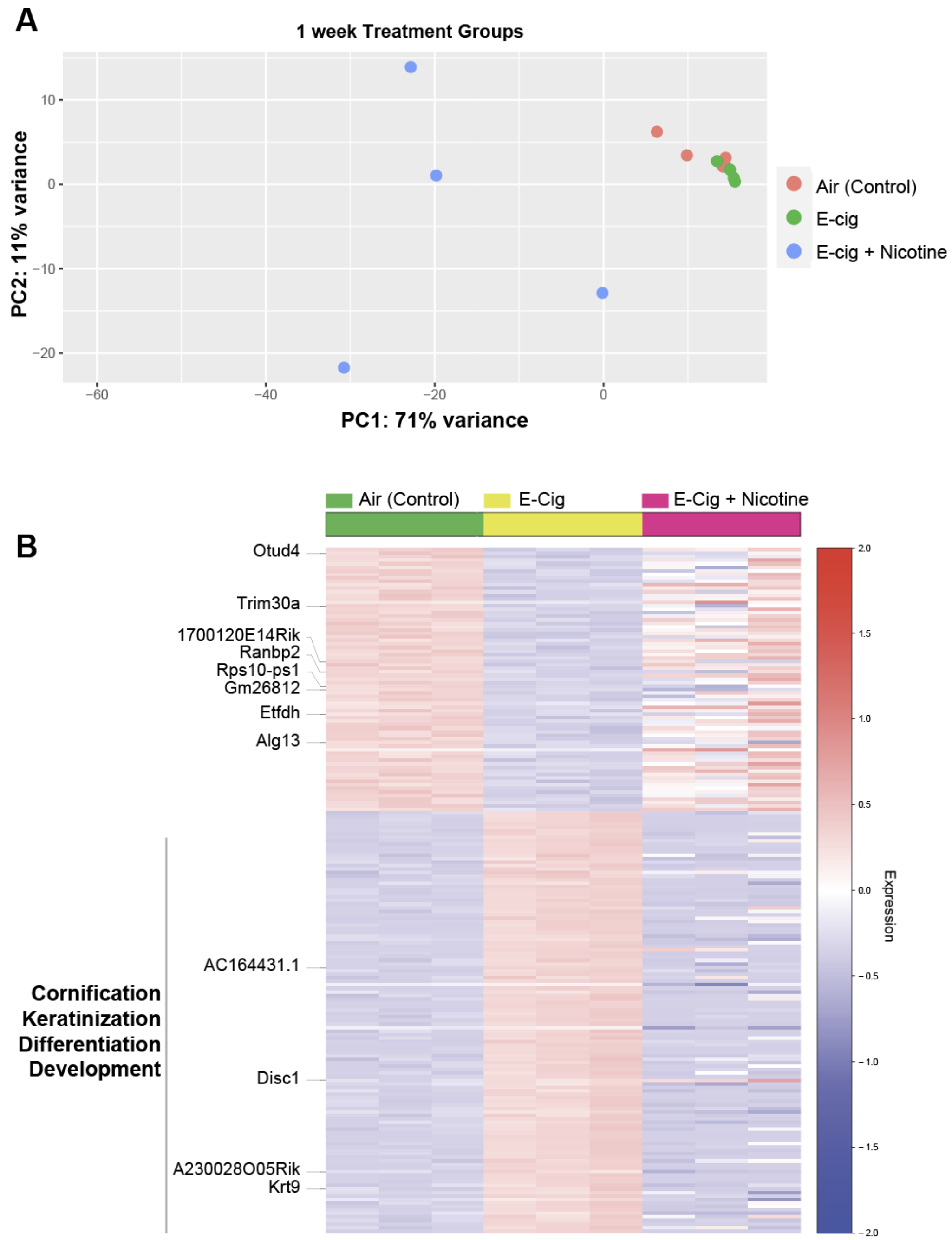

**Figure S2**. **Genes differentially expressed in mouse colon exposed to air or electronic cigarette (ECIG) vapors with and without nicotine**. **A.** Principal component analysis of all 195 (75+120) genes by median absolute deviation. Colon exposed to e-cig vapors (green; n=4), e-cig with nicotine (purple, n =4), or Air (red; n=4). E-cig with nicotine samples differ from air controls and e-cig alone samples along with the first principal component, accounting for 71% of the variability in gene expression. **B.** Heatmap selectively highlighting those genes that are differentially expressed in air control (green) vs. e-cig alone (yellow) groups that are not rescued in the nicotine-containing e-cig group (magenta), as opposed to the large majority of them are indeed returned to levels in the control group (see **Fig 2B**). The upregulated genes support pathways (KEGG analysis) that concern differentiation, cornification and keratinization in the colon.

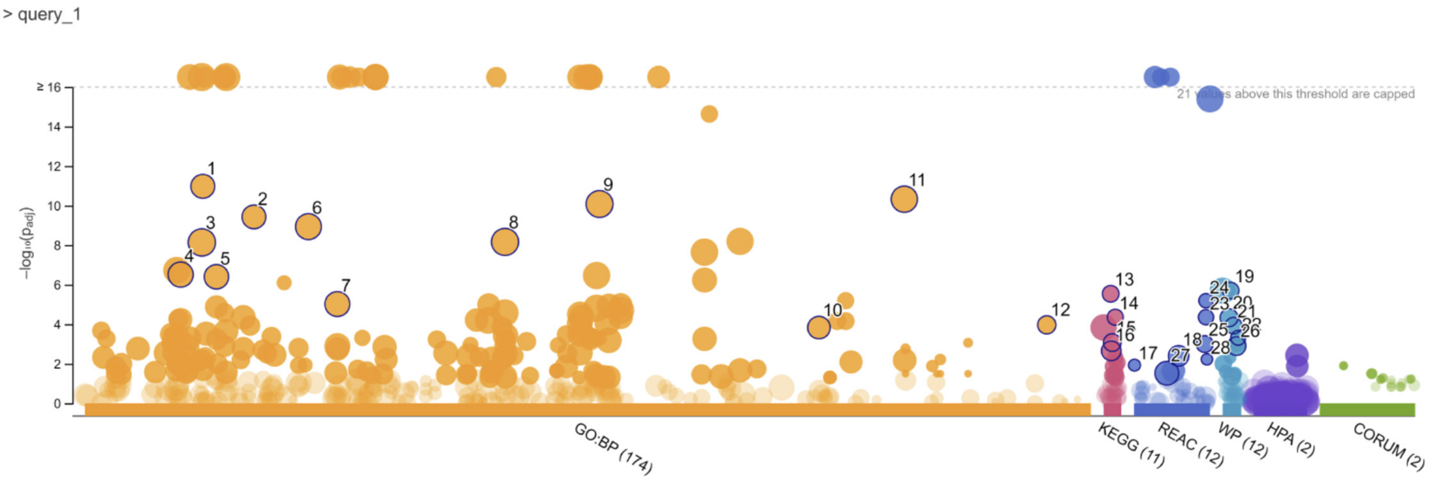

| ID | Source | Term ID | Term Name | $p_{adj}$ (query_1) |
|---|---|---|---|---|
| 1 | GO:BP | GO:0008219 | cell death | $1.060\times10^{-11}$ |
| 2 | GO:BP | GO:0012501 | programmed cell death | $3.723\times10^{-10}$ |
| 3 | GO:BP | GO:0008152 | metabolic process | $7.294\times10^{-9}$ |
| 4 | GO:BP | GO:0006950 | response to stress | $3.077\times10^{-7}$ |
| 5 | GO:BP | GO:0009605 | response to external stimulus | $3.978\times10^{-7}$ |
| 6 | GO:BP | GO:0019538 | protein metabolic process | $1.161\times10^{-9}$ |
| 7 | GO:BP | GO:0023051 | regulation of signaling | $9.560\times10^{-6}$ |
| 8 | GO:BP | GO:0044238 | primary metabolic process | $6.869\times10^{-9}$ |
| 9 | GO:BP | GO:0050896 | response to stimulus | $8.315\times10^{-11}$ |
| 10 | GO:BP | GO:0098542 | defense response to other organism | $1.476\times10^{-4}$ |
| 11 | GO:BP | GO:1901564 | organonitrogen compound metabolic process | $4.735\times10^{-11}$ |
| 12 | GO:BP | GO:1990845 | adaptive thermogenesis | $1.076\times10^{-4}$ |
| 13 | KEGG | KEGG:03320 | PPAR signaling pathway | $2.891\times10^{-6}$ |
| 14 | KEGG | KEGG:04923 | Regulation of lipolysis in adipocytes | $4.425\times10^{-5}$ |
| 15 | KEGG | KEGG:04152 | AMPK signaling pathway | $8.463\times10^{-4}$ |
| 16 | KEGG | KEGG:04060 | Cytokine-cytokine receptor interaction | $2.201\times10^{-3}$ |
| 17 | REAC | REAC:R-HSA-16... | AMPK inhibits chREBP transcriptional activation ac... | $1.171\times10^{-2}$ |
| 18 | REAC | REAC:R-HSA-67... | Neutrophil degranulation | $3.959\times10^{-3}$ |
| 19 | WP | WP:WP3942 | PPAR signaling pathway | $1.976\times10^{-6}$ |
| 20 | WP | WP:WP236 | Adipogenesis | $4.642\times10^{-5}$ |
| 21 | WP | WP:WP1403 | AMP-activated Protein Kinase (AMPK) Signaling | $1.179\times10^{-4}$ |
| 22 | WP | WP:WP3965 | Lipid Metabolism Pathway | $4.876\times10^{-4}$ |
| 23 | REAC | REAC:R-HSA-89... | Triglyceride metabolism | $4.414\times10^{-5}$ |
| 24 | REAC | REAC:R-HSA-16... | Triglyceride catabolism | $6.438\times10^{-6}$ |
| 25 | REAC | REAC:R-HSA-38... | Transcriptional regulation of white adipocyte differ... | $1.004\times10^{-3}$ |
| 26 | WP | WP:WP2877 | Vitamin D Receptor Pathway | $1.275\times10^{-3}$ |
| 27 | REAC | REAC:R-HSA-16... | Immune System | $3.004\times10^{-2}$ |
| 28 | REAC | REAC:R-HSA-89... | VLDL clearance | $5.944\times10^{-3}$ |

**Figure S3**. **Gene ontology (GO) analysis of differentially expressed genes the colons of mice exposed to e-cig compared to air controls.** Manhattan-like plots (top) represent the statistical enrichment (bottom) of pathways were generated using the list of differentially expressed genes and g:GOSt, a core of the g:Profiler that uses statistical enrichment analysis to interpret the functional relevance of gene signatures. The x-axis represents functional terms that are grouped and color-coded by data sources (e.g. GO biological process, orange; KEGG, magenta; Reactome, blue, etc). The y-axis shows the adjusted enrichment p-values in a negative log10 scale. The light circles represent insignificant terms (if available).

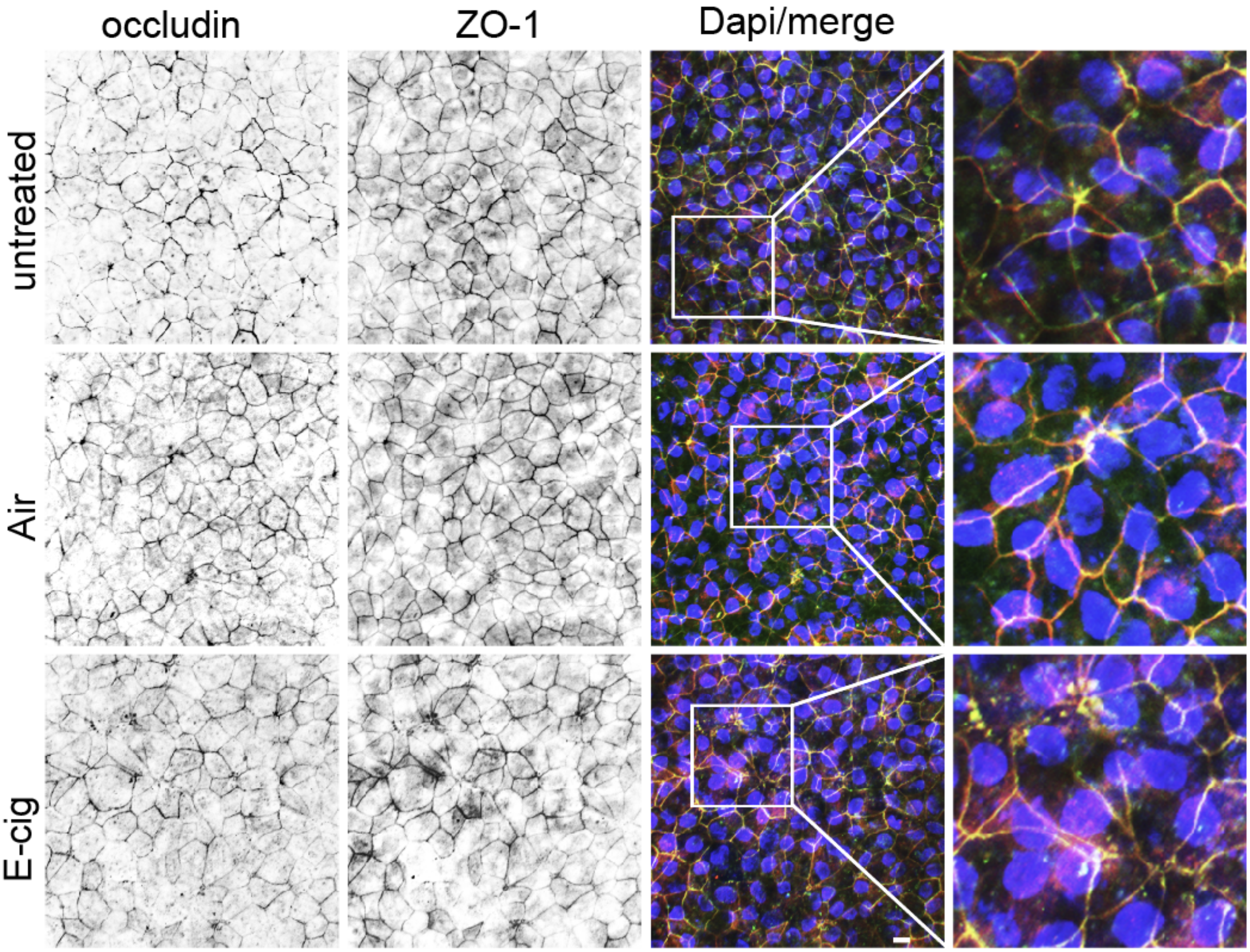

**Figure S4. Confocal microscopic analysis of human EDMs after chronic exposure to e-cig vapor-infused media.** Representative images show Occludin (a TJ marker; green), ZO-1(a TJ marker; red) and DAPI (blue, nuclei) in colonic EDMs either left untreated (top) for 24 h or exposed for the same duration to air (middle) or e-cig (bottom)-vapor-infused media. Scale bar = 10 µm. Cropped images on the right are presented in **Fig 3E**.

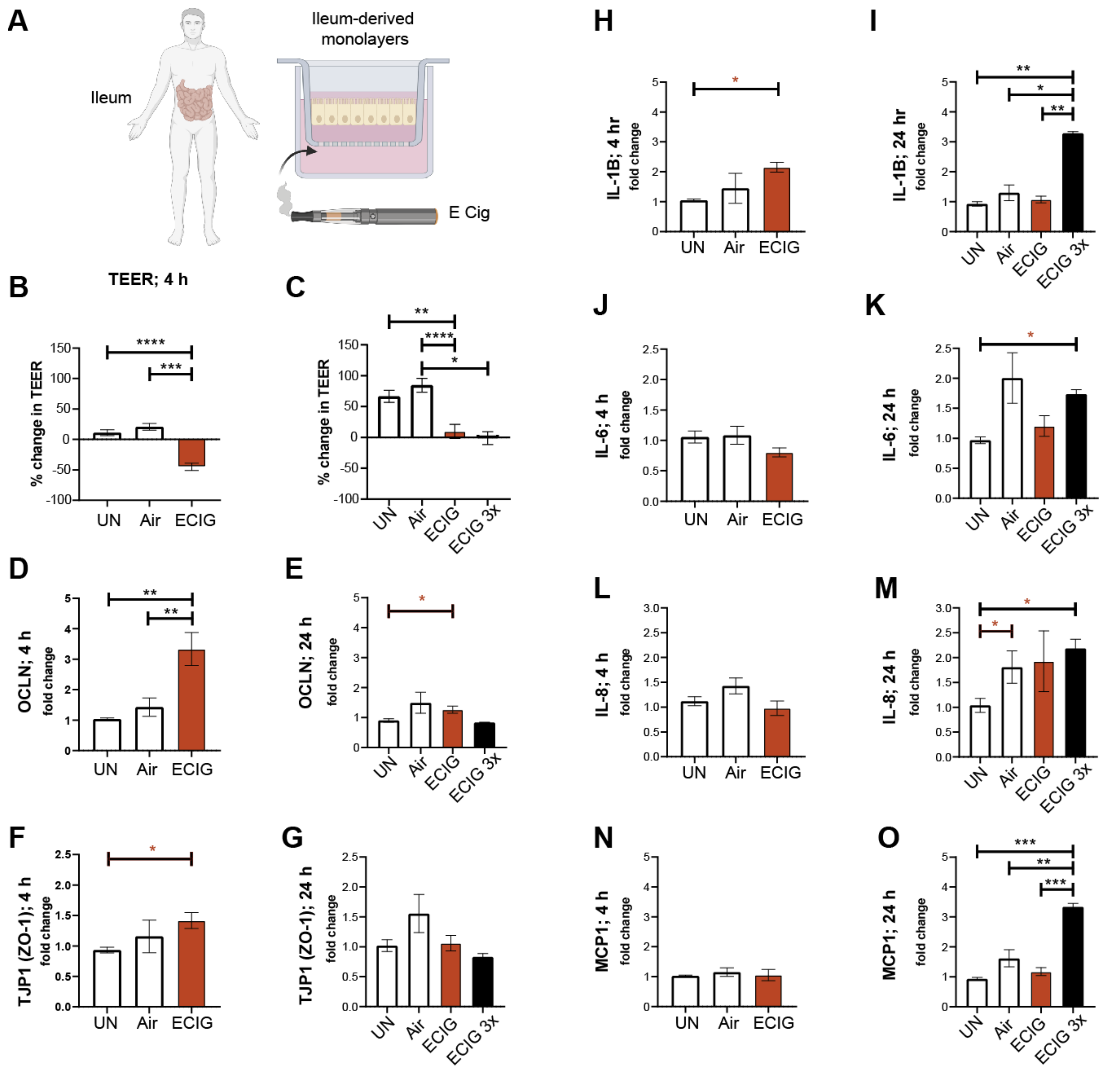

**Figure S5. Nicotine-free e-cig disrupts the human ileal epithelial barrier, triggers inflammation.**

**A.** Schematic displays how enteroids isolated from ileal biopsies of healthy humans were differentiated into polarized enteroid-derived monolayers (EDMs) and exposed to e-cig vapor- infused media on the basolateral side. **B-C**. EDMs were either grown in normal media (UN) or treated with air-infused media (Air) or e-cig vapor-infused media (e-cig) and transepithelial electrical resistance (TEER) was measured over time. The graphs represented the percent change in $\Omega*cm^2$ from three independent experiments and displayed as mean ± SEM. Healthy ileal EDMs were maintained in media or treated with air-infused media or e-cigarette vapor- infused media. After acute exposure of 4 h and chronic exposure of 24 h, mRNA expression was measured after single or multiple exposures. The graphs represented the relative fold change compared to the EDM grown in normal media (UN) and mRNA expression of tight junction markers (**D-G**) and inflammatory cytokines (**H-O**) was measured from at least three independent experiments and displayed as mean ± SEM. One-way ANOVA with Tukey's test and Mann-Whitney (red) test was performed $*p < 0.05$, $**p < 0.01$, $***p < 0.001$ and $****p < 0.0001$.

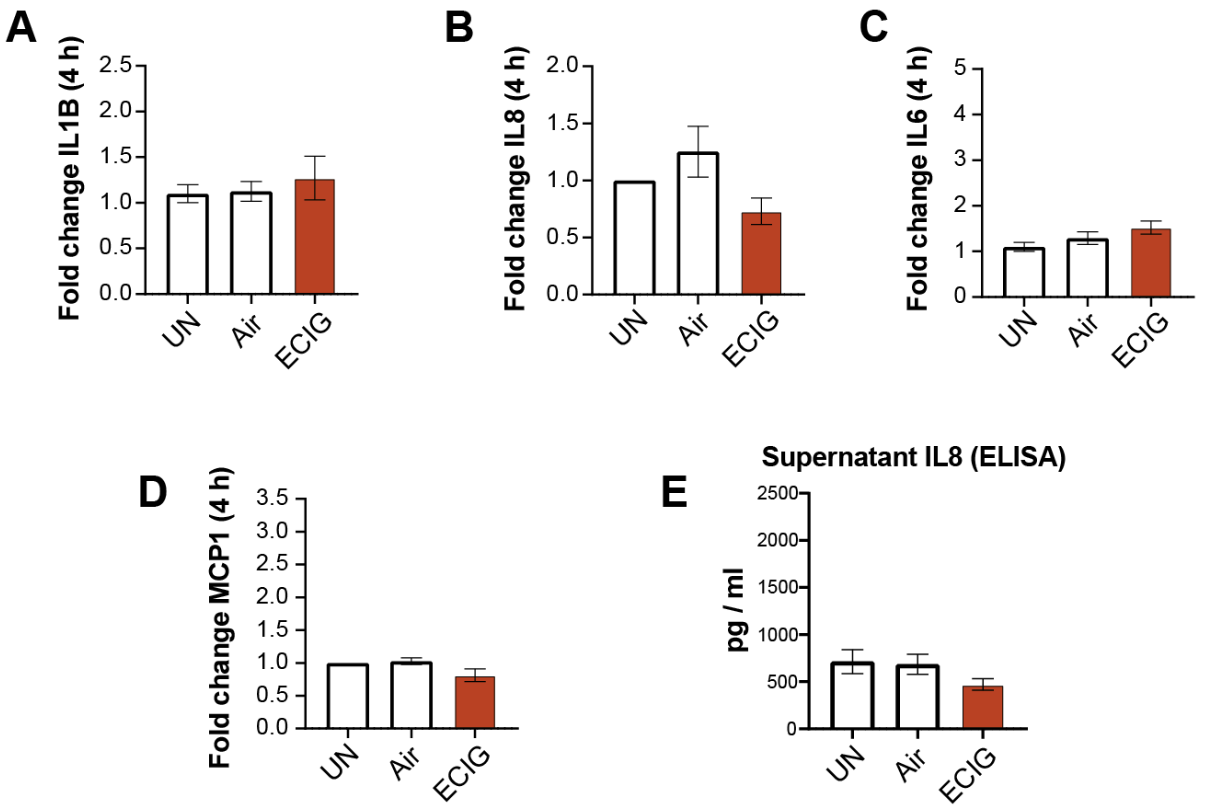

**Figure S6**. **Acute (4 h) exposure to nicotine-free e-cigarettes is not sufficient to trigger colonic epithelial inflammation.** Bar graphs display the relative fold change in mRNA expression of pro-inflammatory cytokines (**A-D**) compared to the EDM grown in normal media (UN) of genes from at least three independent experiments and displayed as mean ± SEM. One-way ANOVA with Tukey's test was performed $*p < 0.05$, $**p < 0.01$ and $***p < 0.001$. (**E)** The supernatant was collected from the basolateral side of EDMs, either grown in normal media (UN) or treated with air-infused media (Air) or e-cigarette vapor- infused media (ECIG) and assessed for pro-inflammatory cytokine IL-8 by ELISA. The graphs represented the concentration of IL-8 from at least three independent experiments and displayed as mean ± SEM.
